# Aphid effectors suppress plant immunity via recruiting defence proteins to processing bodies

**DOI:** 10.1101/2024.11.20.624400

**Authors:** Qun Liu, Anna C. M. Neefjes, Roksolana Kobylinska, Sam T. Mugford, Mar Marzo, James Canham, Mariana Schuster, Renier A. L. van der Hoorn, Yazhou Chen, Saskia A. Hogenhout

**Affiliations:** Department of Crop Genetics, John Innes Centre, Norwich Research Park, Norwich NR4 7UH, UK; Plant Chemetics Laboratory, Department of Biology, University of Oxford, Oxford OX1 3RB, UK; College of Plant Science and Technology, Huazhong Agricultural University, Wuhan 430070, China

## Abstract

Aphids are small insects that have developed specialized mouthparts and effector proteins to establish long-term relationships with plants. The peach-potato aphid, *Myzus persicae*, is a generalist, feeding on many plant species and capable of transmitting numerous pathogens. This study reveals how host-responsive cathepsins B (CathB) in the oral secretions of *M. persicae* facilitate aphid survival by modulating plant immune responses. Aphid CathB localize to processing bodies (p-bodies) and recruit key immune regulators EDS1, PAD4, and ADR1 to these bodies, suppressing plant defenses. A plant protein, Acd28.9 (Hsp20 family), counteracts this CathB activity and contributes to plant resistance to aphids. These findings highlight a novel role for p-bodies in plant immunity and uncover a plant resistance mechanism to aphid infestation.

## Introduction

Aphids are among the most sophisticated feeders in the insect herbivore world. Through over 150 million years of co-evolution with plants (Drohojowska et al., 2020), they have developed specialized mouthparts, known as stylets, which carefully navigate between plant cells to access the phloem (Tjallingii and Esch, 1993). During this process aphids deliver oral secretions (OS), including a variety of effectors, directly into plant cells (Mugford et al., 2016). The effectors help aphids modulate numerous plant processes and establish long-term feeding in the plant vascular tissues (Bos et al., 2010; Snoeck et al., 2022), particularly the nutrient-rich phloem that transports resources from source to sink organs. The success of aphids is evident, as nearly all vascular plants host at least one aphid species, capable of rapidly reproducing, reaching large populations that can cover entire plant tissues. Additionally, aphids are efficient vectors for numerous plant viruses and other pathogens, many of which are heavily reliant on aphids for transmission (Whitfield et al., 2015). Given this success, aphid effectors likely target key plant regulatory processes that not only enable aphids to thrive on plants but also provide a crucial advantage to the pathogens they transmit (Ray and Casteel, 2022). Yet, how these insects manipulate plant processes remains largely unknown.

Among aphids, *M. persicae* is especially intriguing. While most aphid species and insect herbivores have specialized in colonizing one or a few plant species, *M. persicae* is an exceptional generalist, with one of the broadest host ranges among insects (CABI, 2022) — a feat even more remarkable given its predominantly clonal reproduction (Ogawa and Miura, 2014). Previous studies provide evidence that CathB proteins contribute to *M. persicae* ability to colonize plants (Mathers et al., 2017; Chen et al., 2020; Guo et al., 2020). Here, we demonstrate that CathB proteins suppress key plant immunity mechanisms by modulating largely unexplored pathways in plants.

## Results

### CathB proteins in *M. persicae* oral secretions belong to an expanded, host-responsive clade and promote aphid fecundity

Proteomic analysis of *M. persicae* oral secretions (OS) identified CathB proteins, notably CathB6 and CathB12 (Fig. 1A, table S1 and S2). We previously identified these proteins as members of an expanded phylogenetic clade within the CathB gene family of *M. persicae* (Mathers et al., 2017). Genes in this clade exhibit coordinated differential expression depending on the plant host species and are markedly upregulated in aphids feeding on *A. thaliana* (Fig. 1A and fig. S1A) (Mathers et al., 2017; Chen et al., 2020). Aphid CathB proteins consist of three domains: a signal peptide, a prodomain, and a catalytic domain (Aggarwal and Sloane, 2014; Verma et al., 2016) and peptides corresponding to the catalytic domain were detected in aphid OS (Fig. 1B and fig. S1B). Transgenic plants producing the catalytic domains of CathB6, or the closely related CathB3, showed increased aphid progeny, unlike CathB9 transgenic plants (Fig. 1, C and D, fig. S2 and fig. S3). CathB9 is not detected in *M. persicae* OS (Fig. 1A) and lacks a key catalytic triad residue (fig. S4). Interestingly, catalytically inactive CathB3^C116D^ and CathB6^C114D^ lines also supported higher aphid fecundity (Fig. 1C), indicating CathB peptidase activity is not required for this effect.

**Fig. 1.**
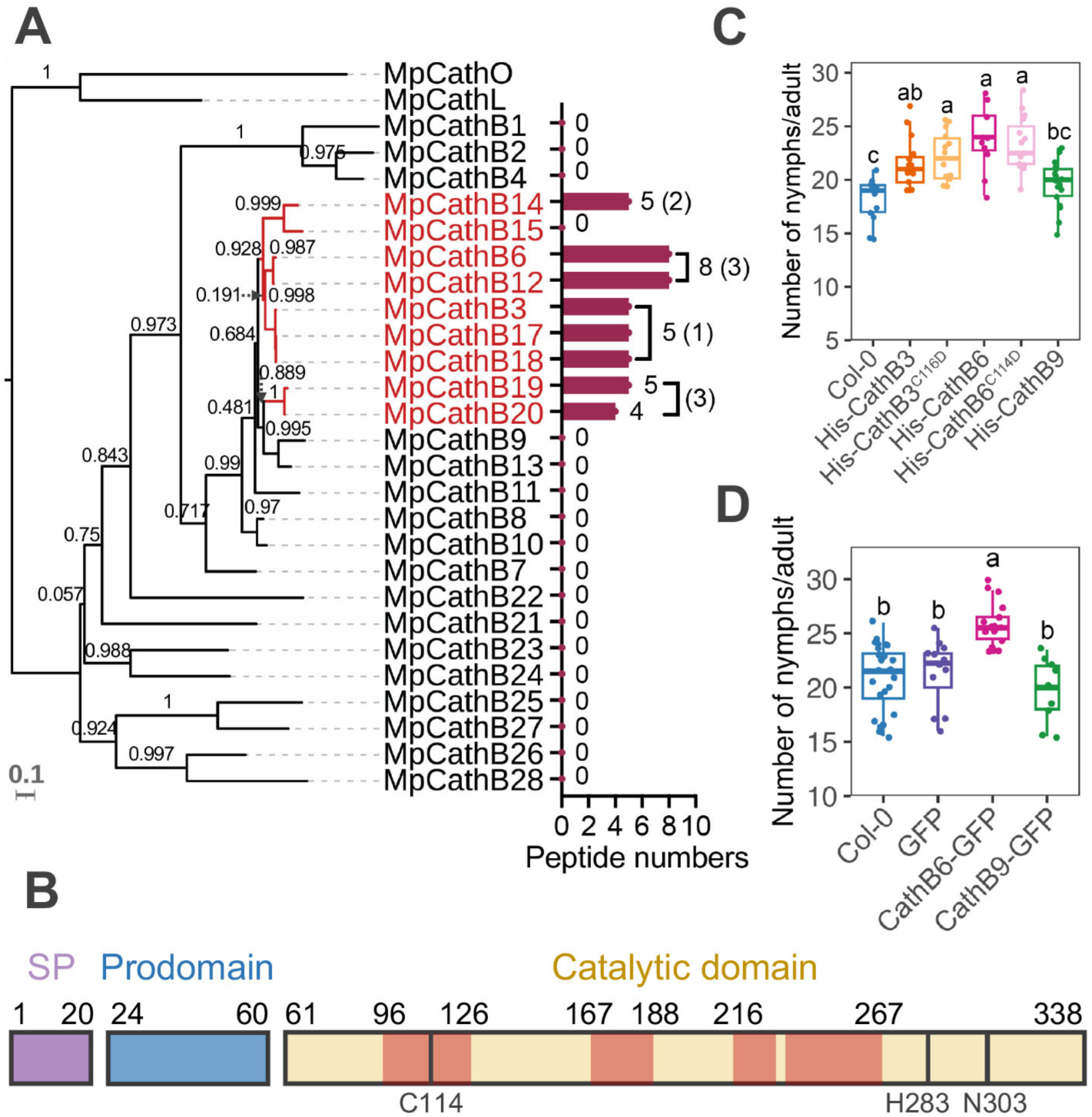
CathB proteins in *M. persicae* oral secretions promote reproduction on ***A. thaliana*. (A**) CathB peptides detected in *M. persicae* oral secretions (OS) match host-responsive proteins showing CathB6 and CathB12 as highly abundant; bars indicate total (unique) peptides. Phylogenetic tree was generated based on alignment of 26 CathB proteins annotated as full-length (table S1) with One CathL and one CathO as outgroups. Scale bar, 0.1 substitutions per site; red branches indicate host-responsive CathBs (fig. S1A). (**B**) Structure of *M. persicae* CathB, with MS-detected regions highlighted in light red (fig. S1B). SP, signal peptide. Catalytic triad residues are marked. First residue of catalytic triad is lacking in CathB9 (fig. S4). (**C** and **D**) Increased fecundity in aphids feeding on plants expressing CathB3, CathB6 or its GFP-tagged version (transgene expression levels, fig. S2A, B). Boxplots show fecundity per adult female (n=9 to 28 replicates per line); letters indicate differences assessed by ANOVA (Tukey method, *p* < 0.05) (additional fecundity assays, fig. S3).

### *M. persicae* CathB localize in processing bodies (p-bodies) in plants

We examined the subcellular localization of CathB6-GFP in *Nicotiana benthamiana* leaves. CathB6-GFP promotes aphid fecundity (Fig. 1D), indicating the GFP fusion does not affect CathB6 function. CathB6-GFP localized to punctate structures of varying sizes, large (> 120 µm²), medium (5 to 120 µm²), and small (< 5 µm²), in the plant cell cytoplasm (Fig. 2A, fig. S5, A and B, and fig. S6A). These puncta are mobile, and smaller puncta often fused with larger ones (Fig. 2B and movie S1), as also evidenced by GFP slowly re-emerging in the larger puncta upon photobleaching (fig. S5, C and D). Similar puncta were observed for CathB6-RFP, CathB6-GFP in the presence of RFP and CathB3-GFP, whereas CathB9-GFP more closely resembled the distribution of free GFP (fig. S7, A and B). CathB6-GFP puncta were also observed in *A. thaliana* protoplasts (fig. S7C).

**Fig. 2.**
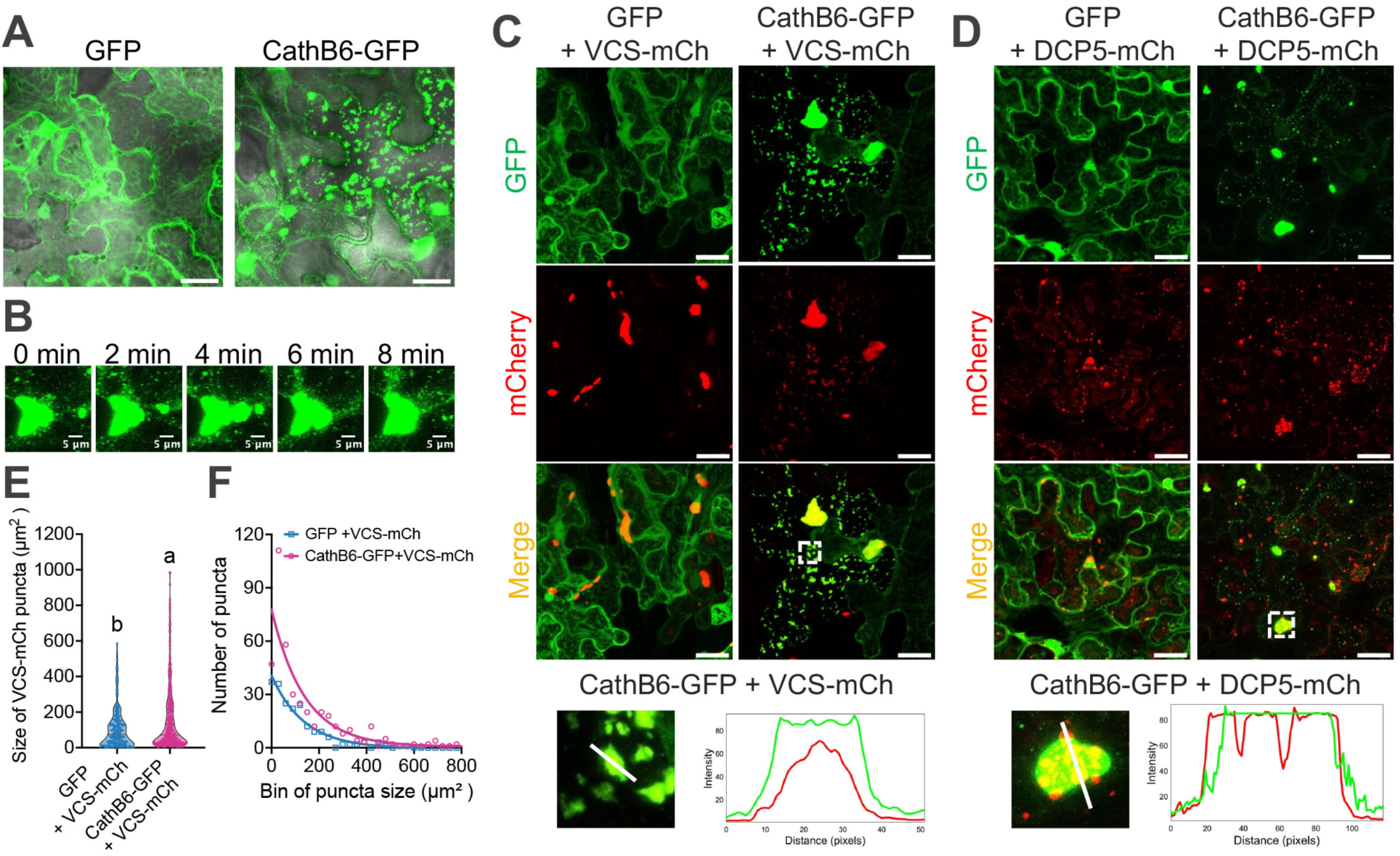
CathB6 forms mobile puncta and colocalizes with processing body markers VCS and DCP5 in *N. benthamiana* leaf cells. **(A**) Confocal images of CathB6-GFP-induced puncta (individual channels, fig. S5A). (**B**) Time-lapse confocal images of a CathB6-GFP puncta fusion (movie S1). (**C** and **D**) Confocal images of CathB6-GFP colocalization with VCS-mCherry (C) and DCP5-mCherry (D), and intensity profiles below the images indicating overlap (more images and puncta size distributions, fig. S8, A and B). Scale bars, 30 μm. (**E** and **F**) Analysis of VCS-mCherry puncta size distributions in the presence of GFP or CathB6-GFP. Significance assessed by two-tailed Student’s t test. Bin size of puncta in (F) is 30 µm².

We co-infiltrated CathB6 constructs with a series of cellular markers that have been reported to form puncta in plant cell cytoplasm (table S3). CathB6-GFP co-localized with the processing body (p-body) scaffolding protein Varicose (VCS) (Fig. 2C, fig. S6B and fig. S8A), decapping protein 5 (DCP5) (Fig. 2D, fig. S6B and fig. S8B) and DCP1 (fig. S8C) (Xu et al., 2006; Steffens et al., 2014; Petre et al., 2016). Whereas VCS-mCherry was depleted in areas occupied by YFP-DCP1 (fig. S8D), CathB6-GFP covered the areas of both VCS-mCherry (Fig. 2C and fig. S8A) and DCP1-RFP (fig. S8C). Similar results were obtained with different fluorescent tags (fig. S9A) and CathB6 also located to p-bodies in *A. thaliana* protoplasts (fig. S9B). CathB6-GFP co-localized at lower frequency with the stress granule marker RBP47b (fig. S10A) (Kosmacz et al., 2018) and did not have obvious colocalizations with autophagosome marker ATG8 (Ibrahim et al., 2023; Pu and Bassham, 2023) and immune regulator NPR1 (Zavaliev et al., 2020) (fig. S10, B and C). Therefore, CathB6 targets p-bodies and partially stress granules in plant cells. P-bodies can convert into stress granules and typically balance the storage, degradation, and translation of mRNAs in cells (Brengues et al., 2005; Kearly et al., 2024). In plants, p-bodies are enriched for mRNAs encoding proteins needed for photomorphogenesis (Jang et al., 2019).

Quantification of puncta size and number shows that CathB6 increases both the number and size of p-bodies (Fig. 2, E and F). In addition, small puncta in these cells co-localize with DCP5-mCherry up to 100% (fig. S8B), and at lower frequencies with VCS and DCP1 (fig. S8, A and C). In contrast, cells with medium and large-sized puncta co-localized mostly with VCS-mCherry and DCP1-RFP (fig. S8, A and C). These data indicate that CathB6 promotes p-body formation, which is shown to occur when translation initiation of mRNAs is blocked (Brengues et al., 2005; Jang et al., 2019). As well, given that DCP5 is one of the first components in p-bodies and recruits VCS and DCP1 (Xu and Chua, 2009; Jang et al., 2019), CathB6-GFP associates with p-bodies early in the p-body formation process.

### CathB6 interacts with EDS1 and sequesters EDS1 and its partner PAD4 to p-bodies

To identify potential plant targets of CathB6, we performed proximity labeling (PL)-mass spectrometry (MS) experiments on stable transgenic *A. thaliana* plants producing CathB6-TurboID fusion. This identified EDS1, with 12 unique peptides and an 8.015-fold enrichment, *p*-value 0.0072, compared to GFP-TurboID controls (fig. S11). EDS1 is a positive regulator of plant basal immunity, effector triggered immunity (ETI) and systemic acquired resistance (SAR), including accumulation of the defense hormone salicylic acid (SA) and *N*-hydroxypipecolic acid (NHP) and associated defense responses (Lapin et al., 2020; Dongus and Parker, 2021; Nair et al., 2021). We found that CathB6 directly interacts with EDS1 in yeast two-hybrid (Y2H) assays (Fig. 3A and fig. S12A), Cath6 pulls down EDS1 from *N. benthamiana* leaves (fig. S13A) and EDS1 pulls down CathB6 from insect cell extracts (fig. S13B). CathB6 interacts with a fragment (EDS1^405-554^) of the EDS1 EP domain (Fig. 3B). CathB3 and CathB6^C114D^ also interact with EDS1 (Fig. 3A and fig. S12A). However, CathB9 did not interact with EDS1 and none of the CathB proteins interacted with PAD4 and SAG101, whereas EDS1 did (Fig. 3A and fig. S12A), as expected (Wagner et al., 2013). This result is consistent with the finding that CathB3, CathB6 and CathB6^C114D^ mutant improved aphid fecundity, while CathB9 did not (Fig. 1, C and D, fig. S2 and fig. S3).

**Fig. 3.**
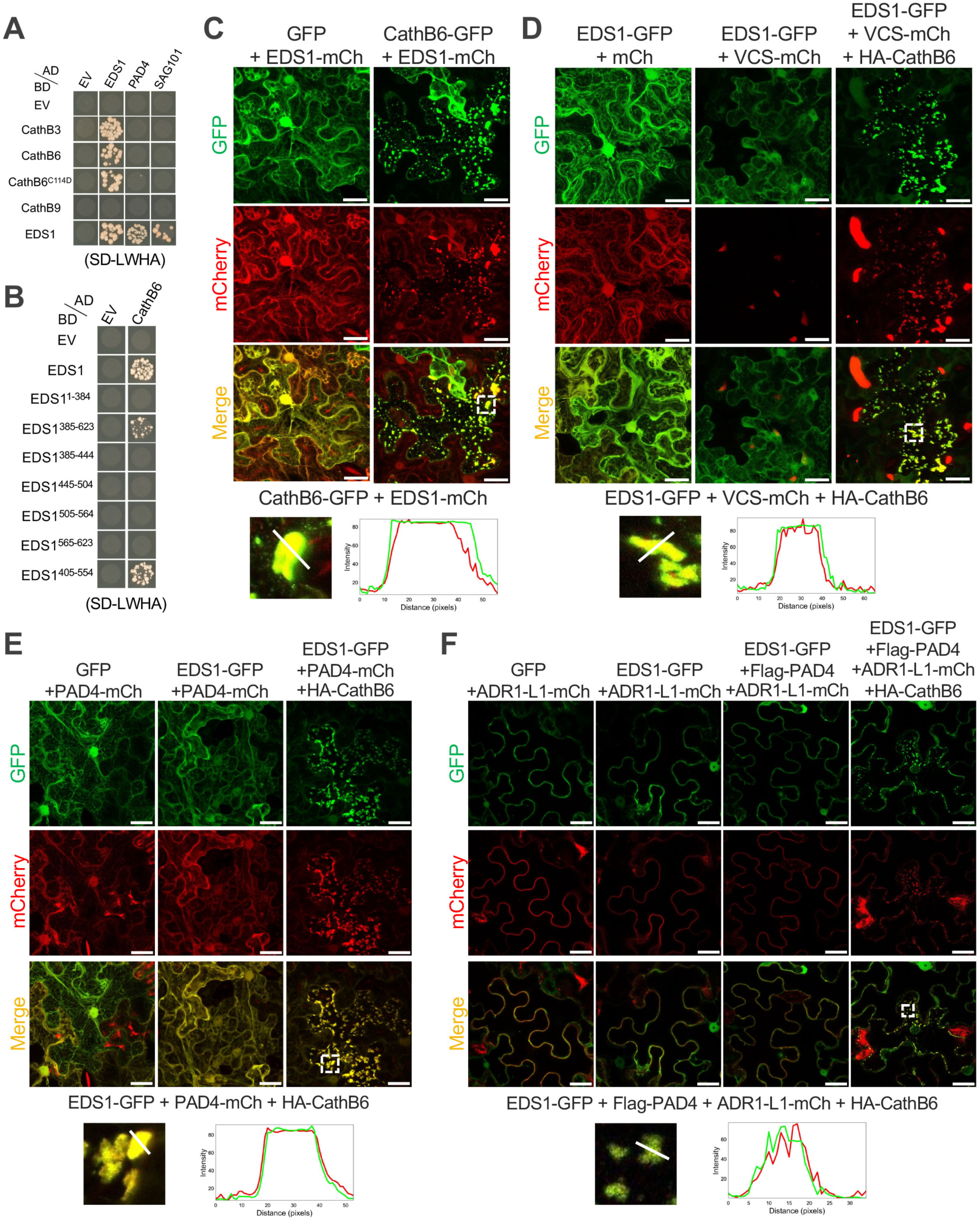
CathB6 interacts with EDS1 and relocates EDS1, PAD4 and ADR1 to p-bodies. (**A** and **B**) Yeast two-hybrid assays on selective SD-LWHA medium (yeast growth on SD-LW medium, fig. S12, A and B). (**C** to **F**) Confocal images illustrating protein localization and colocalization: (C) EDS1-mCherry localized in puncta with CathB6-GFP, (D) colocalization of EDS1-GFP and VCS-mCherry with HA-CathB6, (E) colocalization of EDS1-GFP and PAD4-mCherry in puncta with HA-CathB6, and (F) colocalization of EDS1-GFP and ADR1-L1-mCherry in puncta with Flag-PAD4 and HA-CathB6. Graphs below the confocal images are intensity profiles along the marked lines of the puncta indicated with white boxes in the confocal images above. Scale bars, 30 μm.

Confocal microscopy revealed that in the presence of CathB6-GFP, but not GFP control, EDS1-mCherry forms puncta that co-localize with CathB6-GFP (Fig. 3C and fig. S6C). Similar EDS1-GFP puncta were observed in the presence of CathB6-RFP (fig. S14A) and in *A. thaliana* protoplasts (fig. S14B). Moreover, EDS1-GFP was found to locate in puncta only in the presence HA-CathB6 (fig. S6D and fig. S15A) and these puncta co-localizes with VCS-mCherry (Fig. 3D and fig. S6E), suggesting the puncta are p-bodies. In plants, EDS1 forms heterodimers with PAD4 and SAG101, leading to pathogen restriction or a hypersensitive response (HR)/cell death (Lapin et al., 2020). Both EDS1-GFP and PAD4-mCherry localized to puncta only in the presence of HA-CathB6 (Fig. 3E and fig. S6F). In contrast, EDS1-GFP and SAG101-mCherry localized in nuclei in the absence of HA-CathB6, and these proteins did not form puncta, nor located to other cellular areas, in the presence of HA-CathB6 (fig. S6F and fig. S15B). EDS1 and PAD4 depend on ADR1 for downstream signaling (Saile et al., 2020), and ADR1 also associates with the EDS1 and PAD4 complex (Lapin et al., 2020; Wu et al., 2021). We found that *A. thaliana* ADR1-L1 locates to puncta co-labelled with EDS1-GFP in the presence of Flag-PAD4 and HA-CathB6 (Fig. 3F and fig. S6G). These data indicate that CathB6 sequesters the EDS1-PAD4-ADR1 complex to p-bodies.

### CathB6 modulates *A. thaliana* immunity through EDS1-PAD4 pathway

To assess whether CathB6 influences the expression of genes involved in EDS1-mediated immune signaling, we used the *Pseudomonas fluorescens* Pf0-1 Effector-to-Host Analyzer (EtHAn) system (Thomas et al., 2009) to deliver either the ETI-inducing *P. syringae* effector AvrRps4 or its non-ETI mutant AvrRps4^EEAA^ (Mukhi et al., 2021) into stable *A. thaliana* CathB6-overexpressing lines. AvrRps4, but not AvrRps4 ^EEAA^, induced increased expression of 10 SA- and NHP-regulated genes in wild-type and GFP-expressing control plants, and the induction of all, except EDS1, were significantly suppressed in lines expressing CathB6, including of EDS1 partners PAD4 and SAG101 (Fig. 4A). Similarly, the induction of key marker genes ICS1 and PR1 and a series of other SAR marker genes were profoundly reduced in CathB6 lines (Fig. 4A). These data were consistent across independent repeats (fig. S16). Therefore, CathB6 suppresses the expression of defense genes in the EDS1-mediated response pathway.

**Fig. 4.**
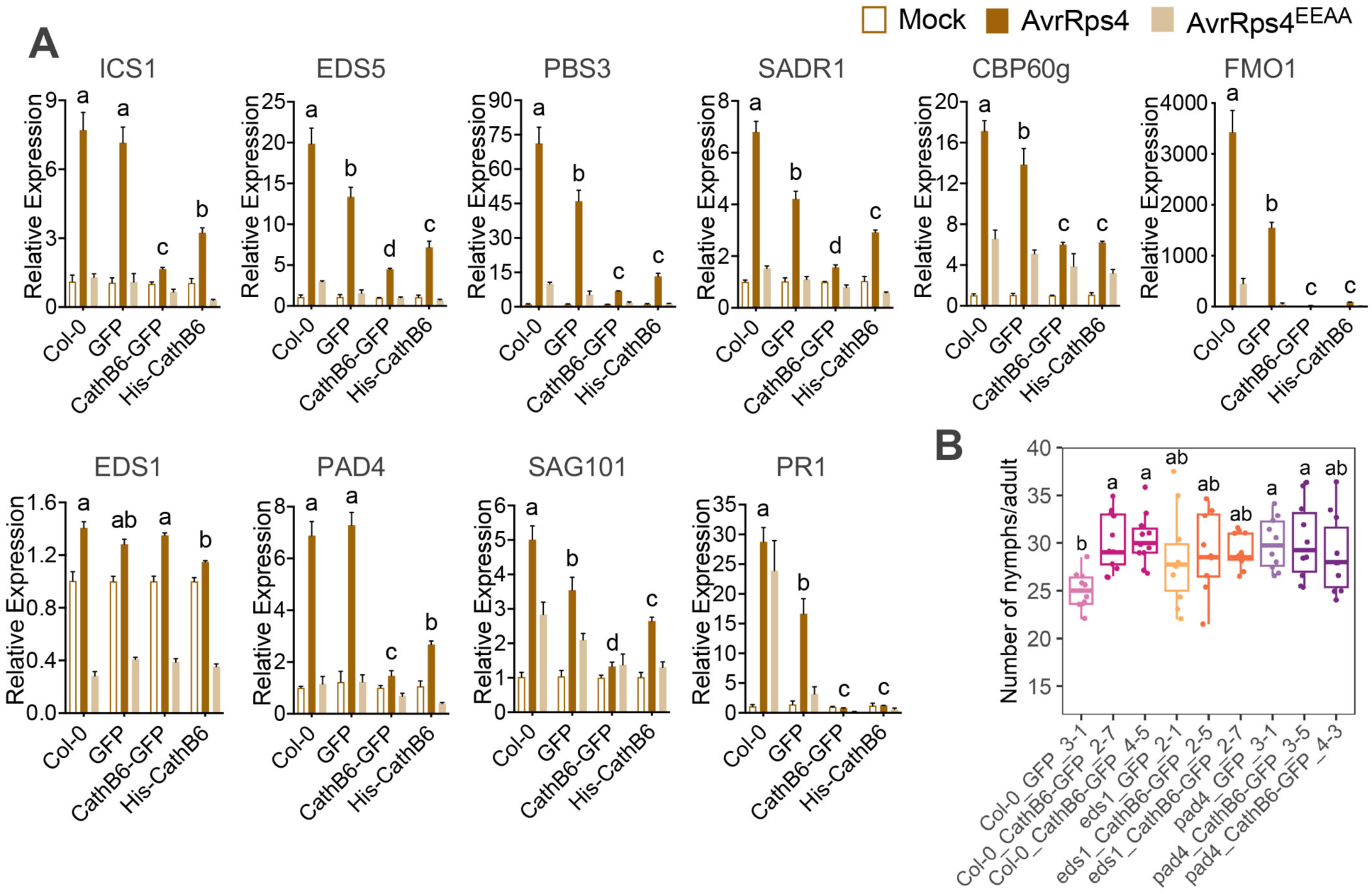
CathB6 inhibits the expression of EDS1-regulated genes involved in the SAR pathway. (**A**) Expression of *A. thaliana* EDS1 and SAR-related genes, including SA biosynthesis (ICS1, EDS5, PBS3) and SAR regulators (PAD4, SAG101, SARD1, CBP60g, FMO1, PR1), in response to mock, Pf0-1 AvrRps4, or Pf0-1 AvrRps4^EEAA^ treatments. Data represent means ± s.e.m. of data of two biological replications; significant differences were analyzed by ANOVA (more independent replicates, fig. S16). (**B**) *M. persicae* fecundity assays on *A. thaliana* lines (transgene expression levels, fig. S2C). Boxplots show fecundity per female aphid (n=9 to12 replicates per line); letters indicate differences determined by ANOVA (Tukey method, *p* < 0.05).

We also investigated whether CathB6 modulates local ETI by analyzing HR levels induced by AvrRps4 in *A. thaliana* Col-0 plants (Mukhi et al., 2021). Controls included *A. thaliana* Col-0 treated with AvrRps4^EEAA^ and *A. thaliana eds1-2* null mutants treated with AvrRps4, both of which should not induce HR responses (Lapin et al., 2019; Mukhi et al., 2021). We observed a slight reduction in HR responses to AvrRps4 in CathB6-GFP lines (14 or 15 out of 20 plants) and His-CathB6 lines (16 or 18 out of 20 plants) compared to wild-type Col-0 (19 out of 20 plants) and GFP plants (19 out of 20 plants). However, the HR response was not completely suppressed (fig. S17). As expected, AvrRps4^EEAA^-treated wild-type plants (0 out of 12 plants) and AvrRps4-treated *eds1-2* null mutants (0 out of 12 plants) showed no HR response (fig. S17). Therefore, CathB6 does not appear to have a clear HR suppression activity.

We generated stable CathB6-GFP transgenic lines in the *eds1-2* and *pad4-1* mutant backgrounds and tested these for aphid fecundity. As before, aphid fecundity was higher on CathB6-GFP in wild-type (Col-0) plants than on GFP-only Col-0 lines (Fig. 4B). Aphid fecundity also increased on *pad4-1* GFP versus wild-type GFP plants and was slightly, but not significantly, higher on *eds1-2* GFP plants compared to wild-type (Fig. 4B). This aligns with reports showing a minor role for EDS1 and a larger effect of PAD4 on aphid fecundity (Dongus et al., 2019; Dongus et al., 2022). Importantly, aphid fecundity was not additionally improved by CathB6 overexpression in *eds1-2* or *pad4-1* mutants, compared to that in Col-0 background (Fig. 4B). These findings indicate that CathB6 modulates plant immunity in the EDS1-PAD4 pathway.

### Hsp20 protein Acd28.9 counteracts CathB-mediated recruitment of EDS1 to p-bodies, and promotes resistance of *A. thaliana* to the aphids

A Hsp20 family protein Acd28.9 was also identified in CathB6 PL-MS (fig. S11). Given that another alpha crystallin (Acd) Hsp20 family protein SLI1 is a resistance factor to *M. persicae* (Kloth et al., 2017; Kloth et al., 2021), we further investigated Acd28.9. We revealed that CathB6 and Acd28.9 interact in Y2H assays (Fig. 5A, fig. S18A), and Cath6 pulls down Acd28.9 from *N. benthamiana* leaves (fig. S18B). The CathB6^266-333^ region interacted with Acd28.9, which differs from the interaction between CathB6 and EDS1, occurring at the CathB6^87-134^ region (fig. S19), and Acd28.9 did not interact with EDS1 (Fig. 5A). Strikingly, CathB6 localization to puncta largely disappeared in the presence of Acd28.9 (Fig. 5B and fig. S6H), indicating that Acd28.9 may counteract CathB6 localization to p-bodies. EDS1-GFP puncta were observed in the presence of HA-CathB6 and free mCherry, while these puncta disappeared in the presence of Acd28.9-mCherry (Fig. 5C and fig. S6I). Therefore, Acd28.9 counteracts CathB6 relocation of EDS1 to p-bodies.

**Fig. 5.**
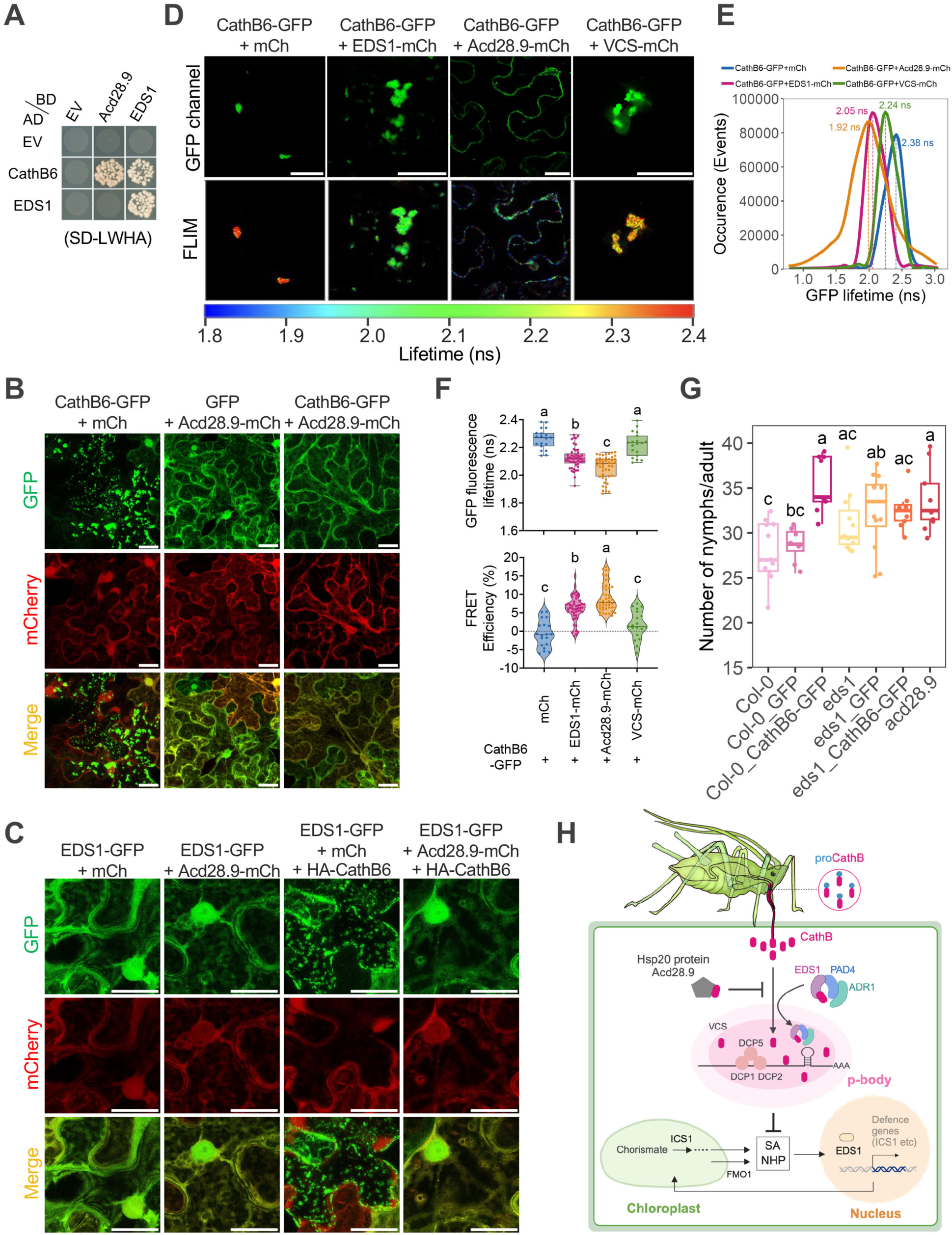
Acd28.9 counteracts CathB6 recruitment of EDS1 to p-bodies and contributes to plant resistance against aphids. (**A**) Yeast two-hybrid assays on selective SD-LWHA medium (yeast growth on SD-LW medium, fig. S12). (**B** and **C**) Confocal images illustrating protein localization and colocalization: Acd28.9-mCherry co-expression depletes CathB6-GFP puncta in *N. benthamiana* (B) and EDS1-GFP forms puncta with HA-CathB6, and does not form puncta with Acd28.9-mCherry and HA-CathB6 (C). (**D** to **F**) FLIM-FRET analysis showing reduced fluorescence lifetime of CathB6-GFP with EDS1 and Acd28.9, but not mCherry and VCS, with CathB6-GFP fluorescence and FLIM images (D), lifetime measurements of CathB6-GFP (E) and statistical lifetime and FRET efficiency mean ± s.e.m (n = 20-49) (F). Scale bars, 30 μm. (**G**) *M. persicae* fecundity assays on *A. thaliana* lines (n=8 to 12 replicates per line). Transgene expression levels, fig. S2C. In (F) and (G), letters indicate differences determined by ANOVA (Tukey method, *p* < 0.05). (**H**) Model: CathB6 recruits EDS1-PAD4-ADR1 to p-bodies, suppressing defence, while Acd28.9 reduces CathB6 localization to p-bodies, counteracting this effect.

To investigate if CathB6 directly interacts with EDS1 and Acd28.9 in plant cells, we conducted Fluorescence Lifetime Imaging Microscopy-based Förster Resonance Energy Transfer (FLIM-FRET) experiments. The lifetime of CathB6-GFP fluorescence was significantly reduced and FRET efficiency significantly improved in the presence of EDS1-mCherry and Acd28.9-mCherry compared to mCherry (Fig. 5, D-F). No reduction of CathB6-GFP fluorescence lifetime was observed in the presence of VCS-mCherry versus mCherry (Fig. 5, D-F), suggesting that though VCS-mCherry was co-localized with CathB6-GFP, it does not directly interact with CathB6-GFP. Thus, CathB6 directly interacts with EDS1 and Acd28.9, but not VCS, within plant cells. Aphid fecundity was increased on *A. thaliana acd28.9* mutant relative to wild-type Col-0, and to a similar level seen on CathB-GFP and *eds1* CathB-GFP lines (Fig. 5G), indicating that Acd28.9 contributes to *A. thaliana* resistance to *M. persicae*.

## Discussion

We report here that CathB6 of the generalist peach-potato aphid *M. persicae* acts as an inhibitor of key plant immunity components. The data supports a model in which CathB6 functions upstream of EDS1, recruiting EDS1, PAD4 and ADR1 to p-bodies, downregulating SAR and promoting aphid fecundity (Fig. 5H). Additionally, CathB6 interacts with a plant Hsp20 protein Acd28.9. Acd28.9 reduces CathB6 and EDS1 localization to p-bodies, and aphid fecundity increases on *acd28.9* mutant lines. Acd28.9 is therefore a resistance protein that counteracts the actions of CathB6 (Fig. 5H).

This study shows that the CathB6 and CathB3 proteins in aphid oral secretions enhance aphid reproduction on *A. thaliana*, unlike CathB9 that is absent from oral secretions. Mutations in the catalytic sites of CathB6 and CathB3 had no impact on aphid fecundity, suggesting that the fecundity-promoting effects of CathB proteins may not rely on their peptidase activity. Therefore, they may function as inactive or “moonlighting” proteases, performing non-catalytic roles, a phenomenon commonly observed in proteins that resemble enzymes (Jeffery, 2020; Werelusz et al., 2024).

We found that CathB6 specifically associates with p-bodies, as indicated by their colocalization with the three p-body markers VCS, DCP1 and DCP5. We observed that CathB6 increases p-body numbers and sizes and that multiple key defense genes are downregulated in the presence CathB. Given that p-bodies enlarge to serve as sites for mRNA storage and promote translational repression (Brengues et al., 2005; Brothers et al., 2023), these data suggest that CathB association with the p-bodies leads to repression of mRNA translation. Moreover, CathB6 recruits EDS1, PAD4 and ADR1 to p-bodies and the genes regulated by EDS1-PAD4-ADR1 module were downregulated. Therefore, CathB appears to target EDS1, PAD4 and ADR1 to p-bodies to downregulate EDS1-PAD4-ADR1 mediated immune pathway.

Our findings are consistent with previous studies showing that PAD4 is involved in aphid defense (Louis et al., 2010) and that aphid fecundity remains unchanged in *pad4 sag101* AtPAD4^R314A^ and *pad4 sag101* AtPAD4^K380A^ plants, where PAD4 EP cavity mutations disrupt EDS1-PAD4 signaling via ADR1 (Dongus et al., 2022). Our data indicate that CathB6 functions upstream of EP domain signaling by recruiting the EDS1-PAD4-ADR1 complex to p-bodies, leading to gene expression downregulation and mitigating the effects of the R314A and K389A mutations. These results also explain the modest increase in aphid fecundity on *eds1-2* plants, as *M. persicae* CathB6 likely suppresses EDS1 directly, while CathB6 indirectly influences PAD4 by downregulating *pad4* expression. The direct inhibition of EDS1, compared to the delayed impact on PAD4, accounts for the higher aphid fecundity observed on *pad4* mutants compared to *eds1* mutants.

CathB expression is upregulated in aphids on *A. thaliana* and *B. rapa* (both Brassicaceae) and downregulated on *N. benthamiana* and potato (both Solanaceae), with expression levels adjusting after host transfer (Mathers et al., 2017; Chen et al., 2020). This is interesting given that EDS1 plays a more minor role in regulating plant defenses in *N. benthamiana* compared to *A. thaliana* (Zönnchen et al., 2022; Huang and Joosten, 2024). Moreover, the EDS1-PAD4 network is more dominant in Brassicaceae, possibly due to a greater diversity of TNLs, while Solanaceae have a higher number of CNLs (Adachi et al., 2019). Nonetheless, the EDS1-PAD4-ADR1 signaling node is broadly present across plants, including monocots, due to its role in both CNL function and basal immunity (Baggs et al., 2020). Aphid CathB expression is upregulated on monocots (fig. S1A), in agreement with a dominant role of the EDS1-PAD4-ADR1 module in regulating immunity in monocots (Lapin et al., 2020). Whether CathB proteins suppress immunity in various plant species is still unknown, though CathB gene duplication, diversification, and positive selection in aphids (Rispe et al., 2008) suggest adaptive immune modulation potential.

Acd28.9, an Hsp20 protein, counteracts CathB6-mediated recruitment of EDS1 to p-bodies, enhancing *A. thaliana* resistance to *M. persicae*. Hsp20 family members are characterized by conserved C-terminal alpha crystallin domains and function as chaperones by preventing protein aggregation (de Jong et al., 1998; Waters and Vierling, 2020). Hsp20 proteins, found across all life domains, generally localize to the cytoplasm but may also occur in organelles and the nucleus, especially in plants (Waters and Vierling, 2020). Plant Hsp20 aid in salt and heat tolerance responses, and some, like RTM2 and SLI1, contribute to virus and insect resistance, respectively (Whitham et al., 2000; Kloth et al., 2021). Thus, at least two Hsp20 proteins contribute to aphid resistance, with Acd28.9 preventing accumulation of CathB6 in plant cell p-bodies.

Relying on plants for all life stages, aphids like *M. persicae* thrive across diverse hosts despite clonal reproduction, underscoring their ability to manipulate plant systems. Additionally, aphid suppression of plant immunity creates an ideal pathway for viruses, facilitating infection and enhancing viral spread. This works shows that the functional study of aphid effectors, especially those in oral secretions, offers valuable insights into plant processes.

## Acknowledgments

We thank Adeline Harant, Sophien Kamoun, Jianhua Huang, Camille-Madeleine Szymansky, Renzo Villena-Gaspar, Joanna Feehan and Jonathan Jones (The Sainsbury Lab), Manuel Gonzalez Fuente and Suayb Üstün (Ruhr-University Bochum) and Tolga Bozkurt (Imperial College London) for providing constructs, seeds and valuable insights. We are grateful to JIC Horticultural Services for growing plants, JIC Entomology Facility for rearing aphids and conducting insect assays, JIC Bioimaging Facility for confocal microscopy, and JIC Proteomics Platform for mass spectrometry analysis.

## Author contributions

Conceptualization: QL, YC, SAH; Funding Acquisition and Project Administration: SAH; Resources: STM, YC, SAH; Supervision: QL, STM, MM, RALH, YC, SAH; Visualization: QL, AN, STM, SAH; Writing – Original Draft: QL, SAH; Writing – Review & Editing: All; Investigation, Methodology, Formal Analysis, and Validation: All.

## Data and materials availability

All data are available in the main text, supplementary materials and public depositories. All generated materials in the manuscript are available upon request.

## Supplementary Materials

### Materials and Methods

#### Aphid colony maintenance

The *M. persicae* Clone O (Fenton et al., 2010) was reared on Chinese cabbage *Brassica rapa* (variety Hilton) or *Arabidopsis thaliana* Col-0 and maintained in a growth chamber (20°C, 14 h light / 10 h dark, 75% humidity) since 2010.

#### Plant growth

Wild-type, mutant or transgenic *A. thaliana* Col-0 plants used for seeds were grown under long day conditions (16 h light/8 h dark) at 20°C with a humidity of 80%. *A. thaliana* Col-0 plants used for fecundity assay were maintained under short day conditions (10 h light/14 h dark) at 22°C with a humidity of 70%. *Nicotiana benthamiana* plants were grown under long day photoperiod (16 h light/8 h dark) at 22°C with a humidity of 80%.

#### Phylogenetic analysis of *M. persicae* cathepsin B (CathB) protein

Genes encoding CathB proteins were extracted from the *M. persicae* clone O v2 genome database (Mathers et al., 2021) and v2.1 annotation (Liu et al., 2024b), identifying a total of 27 CathB proteins, one of which was truncated. A phylogenetic tree was generated using 26 full-length CathB sequences, with cathepsin L (CathL) and cathepsin O (CathO) as outgroups. Full-length protein sequence alignment was performed with MUSCLE algorithm on Phylogeny.fr web server (Dereeper et al., 2008) (https://www.phylogeny.fr/index.cgi). A maximum likelihood phylogenetic tree was constructed in FastTree, using the Shimodaira-Hasegawa test with 1000 resamples (Challa and Neelapu, 2019). The resulting tree was visualized and edited with the Interactive Tree of Life (iTOL) online server (https://itol.embl.de) (Letunic and Bork, 2021).

#### CathB transcript level analysis across nine host plants

The updated CathB annotation was used to analyze the previously published RNA sequencing data of *M. persicae* clone O feeding on nine different plant species were re-analyzed by mapping reads to the v2.1 annotation (Liu et al., 2024b) following similar methods (Chen et al., 2020). Transcript counts per million (TPM) were generated, and differentially expressed CathB genes were identified by comparing transcript levels in aphids feeding on each host species to those in the original colonies on *B. rapa*. Differential expression criteria included *p*-value < 0.05, FDR < 5%, and log2(fold change) > 1. A heatmap was generated, scaled with z-scores (calculated by subtracting the mean expression and dividing by the standard deviation for each CathB gene across all hosts), and integrated with the phylogenetic tree using TBtools (Chen et al., 2023).

#### Generation of CathB transgenic *A. thaliana* plants

The coding sequences corresponding to the catalytic domains of *M. persicae* CathB3 (Lys61-Asn340), CathB6 (Arg61-Asn338) and CathB9 (Glu61-Thr338) were amplified from *M. persicae* clone O cDNA and cloned into vector pBI121 for including a N-terminal His-tag using Gibson assembly cloning methods (Gibson et al., 2009) and to vector pB7FWG2.0 for including a C-terminal GFP-tag using Gateway cloning methods (Liang et al., 2013). After verifying the inserts via sequencing, the constructs were transformed to *Agrobacterium tumefaciens strain* GV3101 and bacterial colonies were grown on LB solid medium containing rifampicin, gentamicin and kanamycin (pBI121-CathB) or spectinomycin (pB7FWG2-CathB) at 28 °C for 24-48 h. Colonies were inoculated into liquid LB medium containing rifampicin, gentamicin and kanamycin or spectinomycin and cultured overnight, followed by plasmid extraction to verify inserts using PCR with gene-specific primers. Positive colonies were then culture at 28 °C in liquid culture and transformed to *A. thaliana* Col-0 plants using the floral dipping method (Bechtold, 1993). Transgenic seeds were harvested and selected on Murashige and Skoog (MS) medium containing 50 μg/mL kanamycin for His-CathB transformants or 20 μg/mL phosphinothricin (BASTA) for CathB-GFP transformants. Seeds from each generation were screened on MS plates with the respective antibiotics to achieve a 3:1 survival-to-death segregation ratio. In T2 or T3 transformed *A. thaliana* plants, a 100% survival rate indicated homozygosity, and these plants were used for fecundity assays.

#### *M. persicae* fecundity assay

Seeds from wild-type, mutant or CathB transgenic *A. thaliana* lines were sown on MS medium containing kanamycin or BASTA. Seedlings from the plates were transplanted to single pots of soil and grown in a controlled environmental room (CER) with a light period of 10 h light/14 h dark at 22°C. Prior to the assay, *M. persicae* clone O adult aphids were transferred from the *A. thaliana* stock colony to three-week-old *A. thaliana* plants, followed by confining the whole plant with a sealed cage. One day later, *M. persicae* nymphs produced by the adult were transferred to different lines of *A. thaliana* plants, with one nymph per plant. The nymphs matured into adults and began reproducing after one week, at which point their offspring were counted on days 7, 9, 11, 13, and 15 post-transfer. Results were calculated as the number of nymphs produced per adult. The assay was conducted three times, representing independent biological replicates, with 8 to 12 plants per line in each replicate. Statistical significance was determined using one-way ANOVA followed by Tukey’s multiple comparison test. Boxplots were generated using R (v4.3.2).

#### Quantification of CathB transgene expression in plants by qRT-PCR

CathB transgenic plants in Col-0, *eds1-2* and *pad4-1* background used for *M. persicae* fecundity assay were quantified for CathB expression using qRT-PCR. In brief, total RNA was isolated from randomly selected plant using RNeasy plant mini kit (QIAGEN) and subsequent DNase treatment using RNase-free DNase I (Thermo Fisher Scientific). cDNA was synthesized from 1 µg of total RNA with RevertAid First Strand cDNA synthesis Kit (Thermo Fisher Scientific). The qRT-PCR reactions were performed on CFX96 Touch Real-Time PCR detection system (BioRad) in triplicate. Each reaction was conducted in 20 µL system containing 10 µL of Maxima SYBR Green master mix (Thermo Fisher Scientific), 0.5 µL of each primer (10 µM), 1 µL of sample cDNA and 8 µL of nuclease-free water (Thermo Fisher Scientific). The cycle conditions were as follows: 95 °C for 10 min, 40 cycles at 95 °C for 15 s and 60 °C for 60 s. All data were normalized to EF1alpha (accession number AT5G60390). Relative quantification was calculated using comparative method as 2^-△Ct^ (Schmittgen and Livak, 2008).

#### Subcellular localization

Plasmids used for transient expression in *N. benthamiana* leaves and transformation of *A. thaliana* protoplasts were generated by amplifying target sequences from cDNA and cloning these into pB7FWG2.0 for adding GFP, pB7RWG2.0 for RFP, or pB7WG2.0 for mCherry tags, using Gibson assembly and Gateway cloning.

For transient expression *N. benthamiana* leaves, the constructs were transformed into *A. tumefaciens* strain GV3101, plated on LB solid medium with appropriate antibiotics, and grown at 28 °C for 24-48 hours. Colonies were picked and verified by PCR with gene-specific primers using plasmid DNA from overnight liquid cultures. Positive colonies were cultured overnight at 28 °C, harvested, and resuspended in infiltration buffer (10 mM MgCl₂, 10 mM MES, pH 5.6) with 100 μM acetosyringone. For infiltration of leaves, GFP- or YFP-tagged constructs were combined with RFP- or mCherry-tagged constructs and pCB301-P19 at an OD600 of 0.3 for each, and infiltrated into the abaxial surface of 4-week-old *N. benthamiana* leaves using a 1 mL needleless syringe. Infiltrations were performed on randomly selected leaves of two different plants. Leaf samples were collected for imaging at 48-72 hours post-infiltration (hpi). Live leaf sections were mounted in water for microscopy.

For *A. thaliana* protoplast transformation, Plasmids were extracted and purified from *E. coli* using QIAGEN Plasmid Midi Kit (QIAGEN). *A. thaliana* mesophyll protoplasts were isolated and transformed as reported (Yoo et al., 2007). Briefly, protoplasts were isolated from leaves of 3-week-old *A. thaliana* plants, which were grown at 22°C under short day (10 h light/14 h dark) conditions with a humidity of 70%. Transformation was conducted by co-transforming 300 μL of fresh protoplast solution (400,000/mL) and 12 μg high quality plasmids of each construct using PEG-calcium method. Transformed protoplasts were kept in dark for 16 h at 22°C. Transformed protoplasts were observed by confocal microscopy the next day.

#### Colocalization analysis using confocal microscopy

Colocalization analyses were conducted with a Leica TCS SP8X upright confocal laser scanning microscope, using either a 20×/0.75 dry or 63×/1.20 water immersion objective (HC PL APO CS). Sequential, unidirectional scans were carried out using the following laser lines: 488 nm, 514 nm (both from 65 mW Argon ion laser), and 580 nm (pulsed SuperK EXTREME supercontinuum white light laser, 470-670 nm, 1.5 mW per line) to excite GFP, YFP, and RFP/mCherry respectively. Fluorescence emissions were collected at 505-540 nm, 529-550 nm, and 595-620 nm. Images were acquired on hybrid detectors at laser power < 5%, with line averaging 2 and a pinhole of 1 Airy unit. Images were generated with varying gains and z-steps. The pixel size was set 0.18 µm × 0.18 µm and pixel dwell time 600 ns.

#### Time-lapse recording using confocal microscopy

Time-lapse experiment was performed with a Leica Stellaris 8 FALCON upright confocal microscope, using a 63×/1.20 water immersion objective (HC PL APO CS2) and xyzt scan mode. Bidirectional scans were carried out using a 20 mW 488 nm laser diode to excite GFP, and fluorescence emissions were collected at 505-540 nm. Images were acquired on hybrid detector S at laser power < 5%. Z-stacks through the entire puncta were taken at minimum intervals. The pixel size was set 0.18 µm × 0.18 µm and pixel dwell time 1.0375 µs.

Image processing and statistical analyses (e.g., puncta size, number, and intensity profiles) were performed with ImageJ (FIJI). Graphs were generated using GraphPad Prism 10 software. Each experiment was independently repeated at least three times.

#### Fluorescence Recovery After Photobleaching (FRAP)

CathB6-GFP constructs were transformed into *A. tumefaciens* strain GV3101 and infiltrated into *N. benthamiana* leaves as described. After 48 hours, leaf sections from infiltrated plants were mounted in water and observed with a Leica Stellaris 8 FALCON upright confocal microscope. Using a 63×/1.20 water immersion objective, a region within a CathB6-GFP punctum was photobleached with 50% laser intensity at 488 nm for three iterations. Recovery was recorded every 2 seconds for a total of 120 seconds post-bleaching. Recovery analysis was performed with FIJI, with fluorescence intensity at each time point normalized using the FRAP profiler v2 plugin. The recovery curve was then analyzed and plotted using GraphPad Prism 10 software.

#### Yeast two-hybrid assay (Y2H)

Coding sequences corresponding to the catalytic domains of *M. persicae* CathB3 (Lys61-Asn340), CathB6 (Arg61-Asn338), CathB9 (Glu61-Thr338), and the CathB6^C114D^ mutant were amplified and cloned into the Gateway vector pDEST-GBKT7 (BD). Full-length sequences of *A. thaliana* EDS1, PAD4, SAG101, and Acd28.9 were amplified and cloned into pDEST-GADT7 (AD). To test interactions between CathB6 and various EDS1 truncations, different fragments of EDS1— including full-length EDS1, the lipase-like domain (EDS1^1-384^), the EP domain (EDS1^385-623^), and additional truncations of the EP domain (EDS1^385-444^, EDS1^445-504^, EDS1^505-564^, EDS1^565-623^ and EDS1^405-554^)—were cloned into pDEST-GBKT7 (BD), with the CathB6 catalytic domain cloned into pDEST-GADT7 (AD) using Gateway cloning methods.

Constructs for protein-protein interaction testing were co-transformed into *Saccharomyces cerevisiae* strain AH109. Empty pDEST-GBKT7 and pDEST-GADT7 vectors served as negative controls. Transformants were first assessed on solid double dropout medium lacking leucine and tryptophan (SD-LW) to confirm the presence of both AD and BD constructs. Interactions between AD and BD fusion proteins were screened on triple dropout medium lacking leucine, tryptophan, and histidine (SD-LWH) supplemented with 5 mM 3-amino-1,2,4-triazole (3-AT) and on quadruple dropout medium lacking leucine, tryptophan, histidine, and adenine (SD-LWHA). Yeast plates were incubated at 28 °C for 5 days before imaging.

#### Coimmunoprecipitation (Co-IP) in *N. benthamiana*

The coding sequences of CathB6 catalytic domain (Arg61-Asn338) and its corresponding catalytic cysteine mutant were constructed to pB7FWG2.0 for CathB6-GFP and CathB6^C114D^-GFP. Full-length coding sequence *A. thaliana* EDS1 or Acd28.9 were amplified and tagged with 3×HA at N-terminal, then ligated into Gateway vector pB7WG2.0. Constructs were separately transformed to *A. tumefaciens* strain GV3101 and co-infiltrated into 4-week-old *N. benthamiana* leaves using same methods above. Infiltrated leaves were harvested between 48-72 hpi. Total proteins were extracted with extraction buffer [150 mM Tris-HCl (pH 7.5), 150 mM NaCl, 10 mM EDTA, 10% Glycerol, 20 µM NaF, 10 mM DTT, 0.5% (w/v) PVPP, 1% protease Inhibitor cocktail (Sigma), 0.2% Igepal]. GFPtrap beads (Chromotech) were then added into the extracts and incubated on a rotor wheel at 4 °C overnight. The next day, beads were washed with washing buffer [10 mM Tris-HCl (pH 7.5), 150 mM NaCl, 0.5 mM EDTA, 0.2% Igepal] for six times. After washing, proteins were eluted from the beads using 4 × LDS Sample Loading Buffer with 10 mM DTT, then loaded onto 12% SDS-PAGE gels (Invitrogen) and transferred to 0.45 μm PVDF membranes. The membranes were then probed with anti-GFP and anti-HA antibodies.

#### Protein expression in Sf9 cells using the baculovirus expression system

The coding sequence of the CathB6 catalytic domain (Arg61-Asn338) and full-length *A. thaliana* EDS1 were individually tagged with GFP and HA, then cloned into the pFastBac HTB vector using Gibson assembly. After sequencing confirmation, the constructs were transformed into *E. coli* DH10Bac cells, which contain a baculovirus shuttle vector—a bacterial artificial chromosome (BAC) carrying the full genome of *Autographa californica* multiple-nucleocapsid nucleopolyhedrosis virus (AcMNPV).

Following transformation, cells were plated on LB agar with 50 µg/mL kanamycin, 10 µg/mL tetracycline, 7 µg/mL gentamicin, 40 µg/mL IPTG, and 100 µg/mL X-Gal. White colonies were selected, streaked on fresh plates, and screened through four cycles. Bacmids were extracted from positive colonies using an SDS lysis, chloroform extraction, and ethanol precipitation method. (McCarthy and Romanowski, 2008). After verifying recombinant bacmids by PCR, they were transfected into Sf9 cells with Lipofectamine 2000 (Thermo Fisher Scientific).

Five to six days post-transfection, when cells exhibited signs of infection—such as increased size, granularity, cessation of growth, detachment, and lysis—the cell medium containing viral particles was collected as the P0 viral stock. This stock was used to infect fresh Sf9 cells, generating a P1 viral stock. The virus was further amplified through two additional cycles, producing P3 viral stock for optimal protein expression.

#### In vitro co-immunoprecipitation (co-IP) following expression in Sf9 cells

P3 viral stocks of CathB6-GFP and HA-EDS1 were used to infect fresh Sf9 cells for protein expression. Cells were harvested 48-72 hours post-infection, and total protein was extracted using a lysis buffer [25 mM Tris-HCl (pH 7.4), 150 mM NaCl, 1% NP-40, 1 mM EDTA, 5% glycerol] with or without 1% protease inhibitor cocktail (Sigma). Anti-HA magnetic beads (Thermo Fisher Scientific) were added to the extracts, followed by overnight incubation on a rotating wheel at 4 °C. The following day, the beads were washed six times with washing buffer [Tris-buffered saline (TBST) with 0.05% Tween-20]. After washing, proteins were eluted from the beads using 4 × LDS Sample Loading Buffer with 10 mM DTT, then loaded onto 12% SDS-PAGE gels (Invitrogen) and transferred to 0.45 μm PVDF membranes. The membranes were then probed with anti-GFP, anti-CathB6, and anti-EDS1 antibodies.

#### Expression analysis of EDS1-regulated genes by qRT-PCR

The *Pseudomonas fluorescens* Pf0-1(T3SS) EtHAn (Effector-to-Host Analyzer) system (Thomas et al., 2009) carrying AvrRps4 or AvrRps4^EEAA^ strains (Mukhi et al., 2021) was grown on King’s B agar plate supplemented with 30 µg/mL Chloramphenicol and 50 µg/mL Genetamycin at 28 °C overnight. Bacteria were harvested, resuspended in infiltration buffer (100 mM MES, 100 mM MgCl₂), and adjusted to an OD600 of 0.2. The bacterial suspensions were infiltrated into the abaxial surface of 5-week-old *A. thaliana* plants using a 1 mL needleless syringe. Three plants from each *A. thaliana* line were randomly selected, with one leaf per plant infiltrated with Pf0-1 AvrRps4 and another with AvrRps4^EEAA^. Mock samples were infiltrated with buffer only.

Leaves were harvested 8 hours post-infiltration (hpi), and total RNA was isolated using the RNeasy Plant Mini Kit (QIAGEN), followed by DNase treatment with RNase-free DNase I (Thermo Fisher Scientific). cDNA was synthesized from 1 µg of total RNA using the RevertAid First Strand cDNA Synthesis Kit (Thermo Fisher Scientific). qRT-PCR reactions were performed in triplicate on a CFX96 Touch Real-Time PCR Detection System (BioRad) in a 20 µL reaction volume containing 10 µL of Maxima SYBR Green master mix (Thermo Fisher Scientific), 0.5 µL of each primer (10 µM), 1 µL of cDNA, and 8 µL of nuclease-free water (Thermo Fisher Scientific). Cycling conditions were 95 °C for 10 min, followed by 40 cycles of 95 °C for 15 s and 60 °C for 60 s.

All data were normalized to EF1alpha (accession number AT5G60390), and relative expression was calculated using the 2^-ΔΔCt^ method (Schmittgen and Livak, 2008), with mock samples as the reference.

#### Hypersensitive Response (HR) assay in Arabidopsis

The *Pseudomonas fluorescens* Pf0-1EtHAn system carrying either the AvrRps4 or AvrRps4^EEAA^ strain was prepared and infiltrated into 5-week-old *A. thaliana* plants as described. Infiltrations were conducted on randomly selected leaves, with each plant receiving one infiltration of Pf0-1 AvrRps4 on one leaf and AvrRps4^EEAA^ on another. A total of 12 to 20 plants were treated. Plants were then observed and assessed for cell death as an indicator of hypersensitive response 24 hours post-infiltration.

#### FLIM-FRET analysis

Constructs expressing genes of interest (CathB6-GFP, mCherry, EDS1-mCherry, Acd28.9-mCherry, and VCS-mCherry) were transformed into *A. tumefaciens* strain GV3101 and co-infiltrated into 4-week-old *N. benthamiana* leaves as described. After 48 hours, leaf sections were examined using a Leica Stellaris 8 FALCON scanning confocal microscope with a 63×/1.20 water immersion objective. FLIM experiments were conducted in TCSPC (Time-Correlated Single Photon Counting) mode with highly sensitive photon detectors (HyD X). GFP was excited using a pulsed white light laser (WLL) at 488 nm, with emission collected between 505-520 nm. Laser power was maintained below 1% to avoid sample bleaching, with a frequency set to 80 MHz.

The instrument response function (IRF) was calibrated using potassium iodide (KI) and Erythrosine B, as previously described (67). FLIM data sets were recorded using Leica LAS X software with the FLIM Wizard, with each image acquisition continued until reaching a minimum of 5000 photon counts per pixel. Images were acquired at 128 × 128 pixels resolution with a pixel dwell time of 19 µs, and laser power was adjusted to a maximum count rate of 2000 kcounts per second. Each acquisition was stopped after 40 frames.

Data analysis involved measuring the excited-state lifetime values of regions of interest (ROIs), with calculations performed using LAS X software. Lifetime values were obtained by re-convolution fitting, selecting a two-exponential fit, and ensuring a *χ*^2^ value between 0.90 and 1.20. The amplitude-weighted mean lifetime (τ) was used for comparison, and FRET efficiency was calculated by comparing the lifetime of CathB6-GFP co-expressed with EDS1-mCherry, Acd28.9-mCherry, and VCS-mCherry (τₓ) to that of CathB6-GFP with mCherry alone (τ₀) using the formula: FRET efficiency = 1 - (τ_ₓ_ / τ_₀_).

Boxplots and violin plots were generated using GraphPad Prism 10, and statistical significance was assessed by one-way ANOVA followed by Tukey’s multiple comparison test.

---

**Fig. S1.**
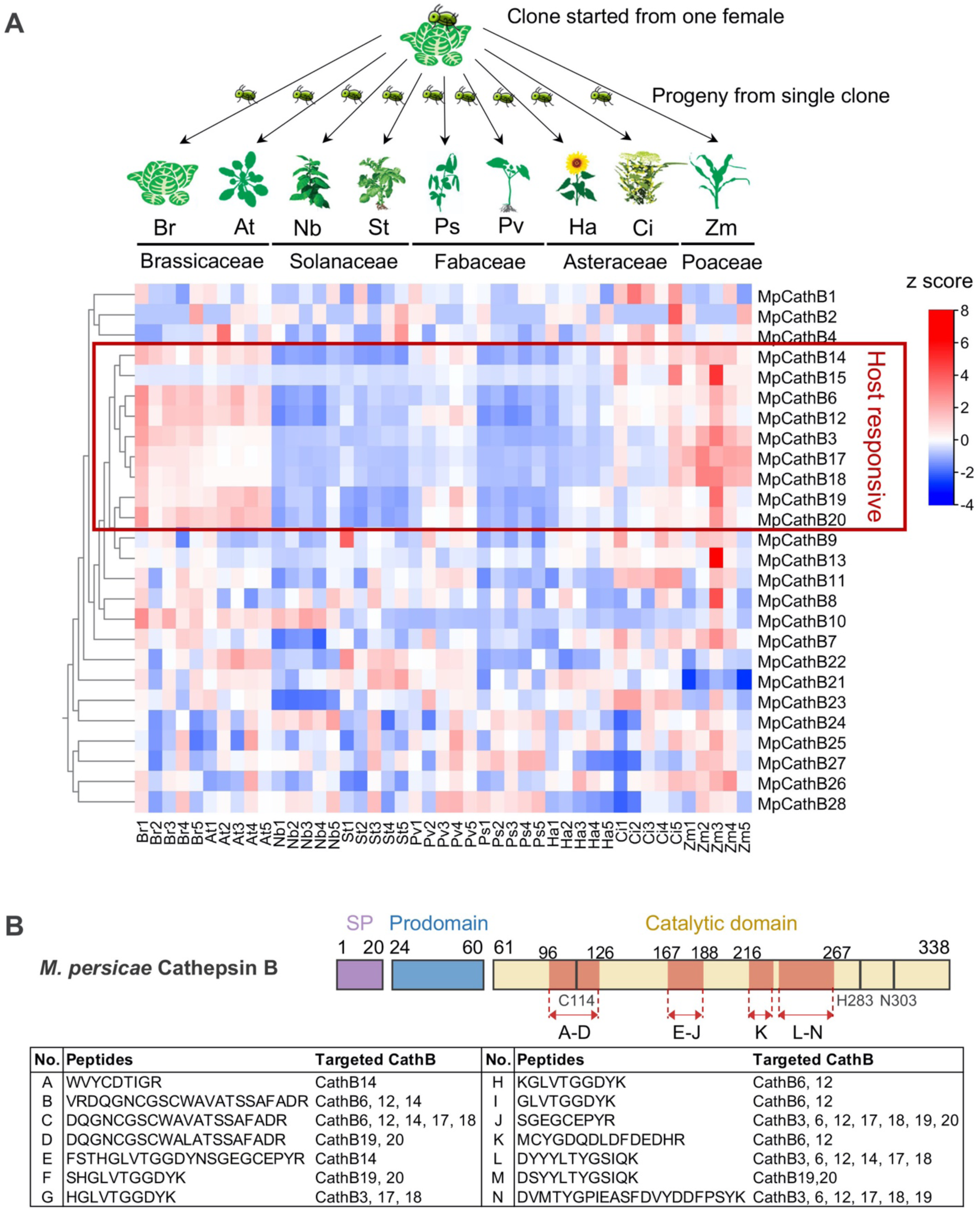
*M. persicae* CathB genes are host responsive and present in oral secretion (OS). (**A**) *M. persicae* CathB differentially express in the aphids on different host plants. (Top) Schematic overview of *M. persicae* clonal populations transferred to 9 plant species from 5 families, as shown. Br, *Brassica rapa*; At, *A. thaliana*; Nb, *Nicotiana benthamiana*; St, *Solanum tuberosum*; Ps, *Pisum sativum*; Pv, *Phaseolus vulgaris* (Pv); Ha, *Helianthus annuus*; Ci, *Chrysanthemum indicum*; Zm, *Zea mays* (Zm). (Bottom) Heatmap of CathB expression values (TPM) of *M. persicae* on nine plant species at five biological replicates per host plant (as indicated below the heatmap). CathB marked in a red rectangle were found to have synchronized up- or down regulation depending on the plant species the aphids were reared on. (**B**) Location of CathB peptides detected in *M. persicae* oral secretions (OS). The peptides matched the CathB catalytic domain. The signal peptide (SP) and prodomain are also indicated. Detected peptides by MS are marked in light red. The catalytic triad residues (Cys114, His283 and Asn303) are marked. Table shows the specific peptides that were detected and which CathB they are likely derived from. The A to N numbering match those in the schematic overview above. Full proteome dataset of *M. persicae* OS is shown in (Liu et al., 2024b).

**Fig. S2.**
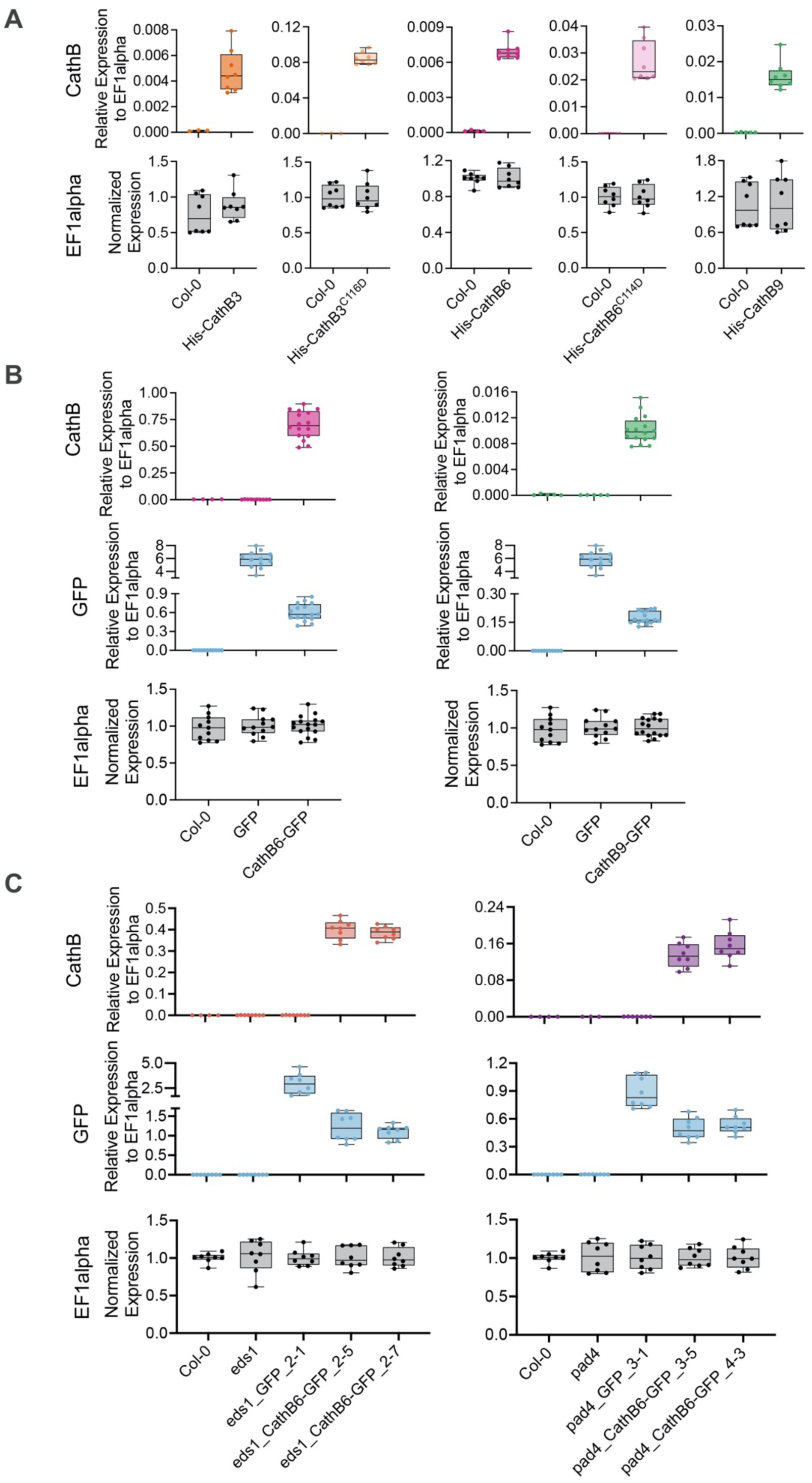
Quantification of transgene expression levels in CathB3, B6 and B9 stable transgenic *A. thaliana* lines used for *M. persicae* fecundity assay. (**A** to **C**) Quantifications of transgene transcript levels by qRT-PCR using specific primers to CathB3 (and B3^C116D^), B6 (and B6^C114D^) and B9 (A) and CathB6 and GFP (B and C). Samples for transgene quantification were collected from three single plants and quantified with n = 3 to 12 repeats (dots on boxplots). Relative transcript levels were determined by qRT-PCR and normalized to the internal control EF1alpha. This figure relates to Figures 1C, 1D and 4B.

**Fig. S3.**
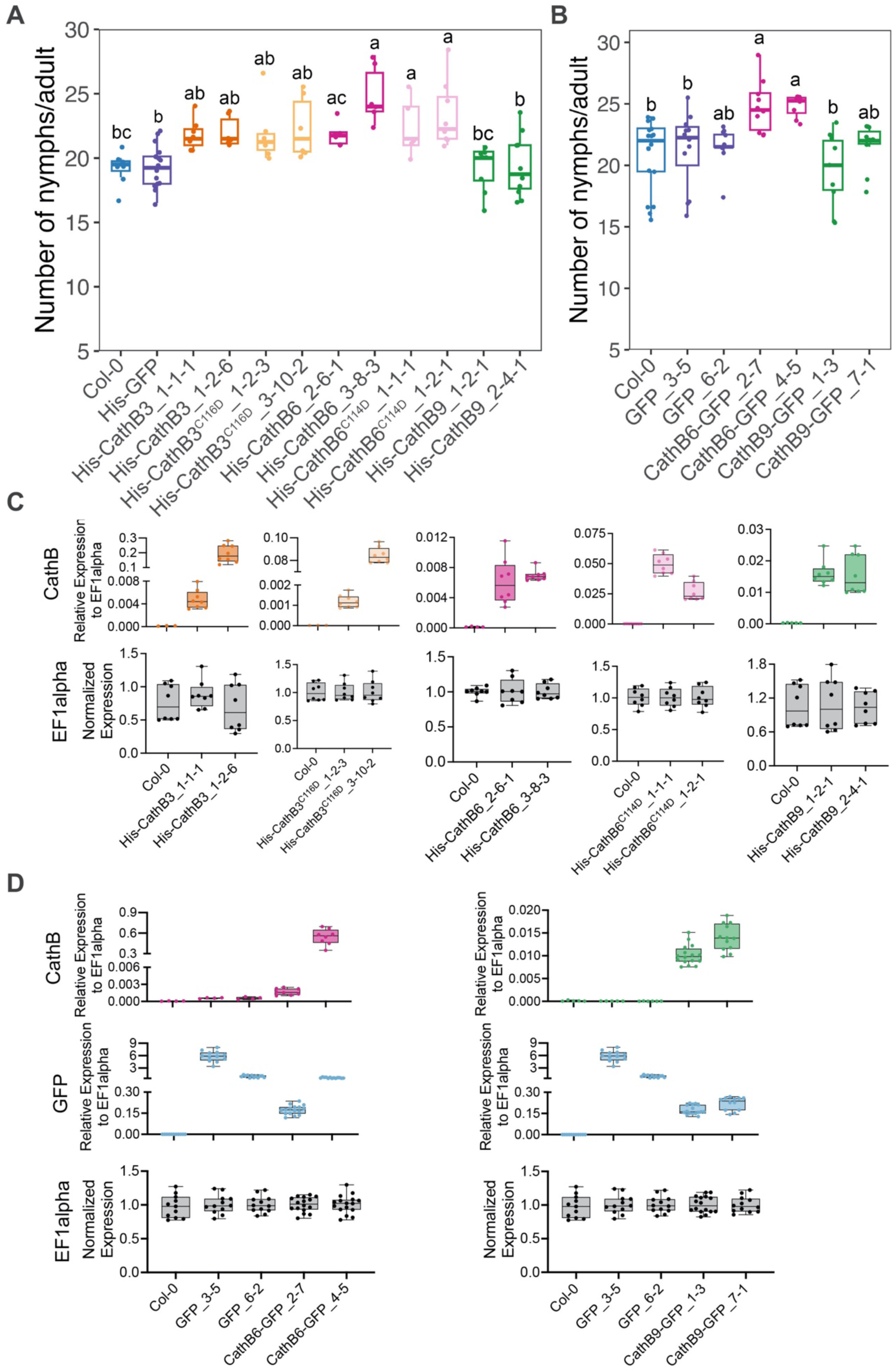
*M. persicae* reproduction is enhanced on transgenic plants stably expressing CathB3, CathB6, or their catalytic mutants, but not on plants expressing CathB9. (**A** and **B**) Aphid fecundity assays on stable His-GFP, His-CathB3, His-CathB3^C116D^, His-CathB6, His-CathB6^C114D^, and His-CathB9 (A) and GFP, CathB6-GFP and CathB9-GFP (B) transgenic *A. thaliana* lines. Box plots show the distribution of nymphs produced by an adult female aphid per plant (dots) collected from n = 6 to 16 female aphids per *A. thaliana* line. Statistical significance was determined by ANOVA (Tukey method, *p* < 0.05) and presented as different letters. (**C** and **D**) Quantifications of transgene transcript levels by qRT-PCR using specific primers to CathB3 (and B3^C116D^), B6 (and B6^C114D^) and B9 (C) and CathB6 and GFP (D). Samples for transgene quantification were collected from three single plants and quantified with n = 3 to 12 repeats (dots on boxplots). Quantification refers to relative transcript levels which were determined by qRT-PCR and normalized to the internal control EF1alpha.

**Fig. S4.**
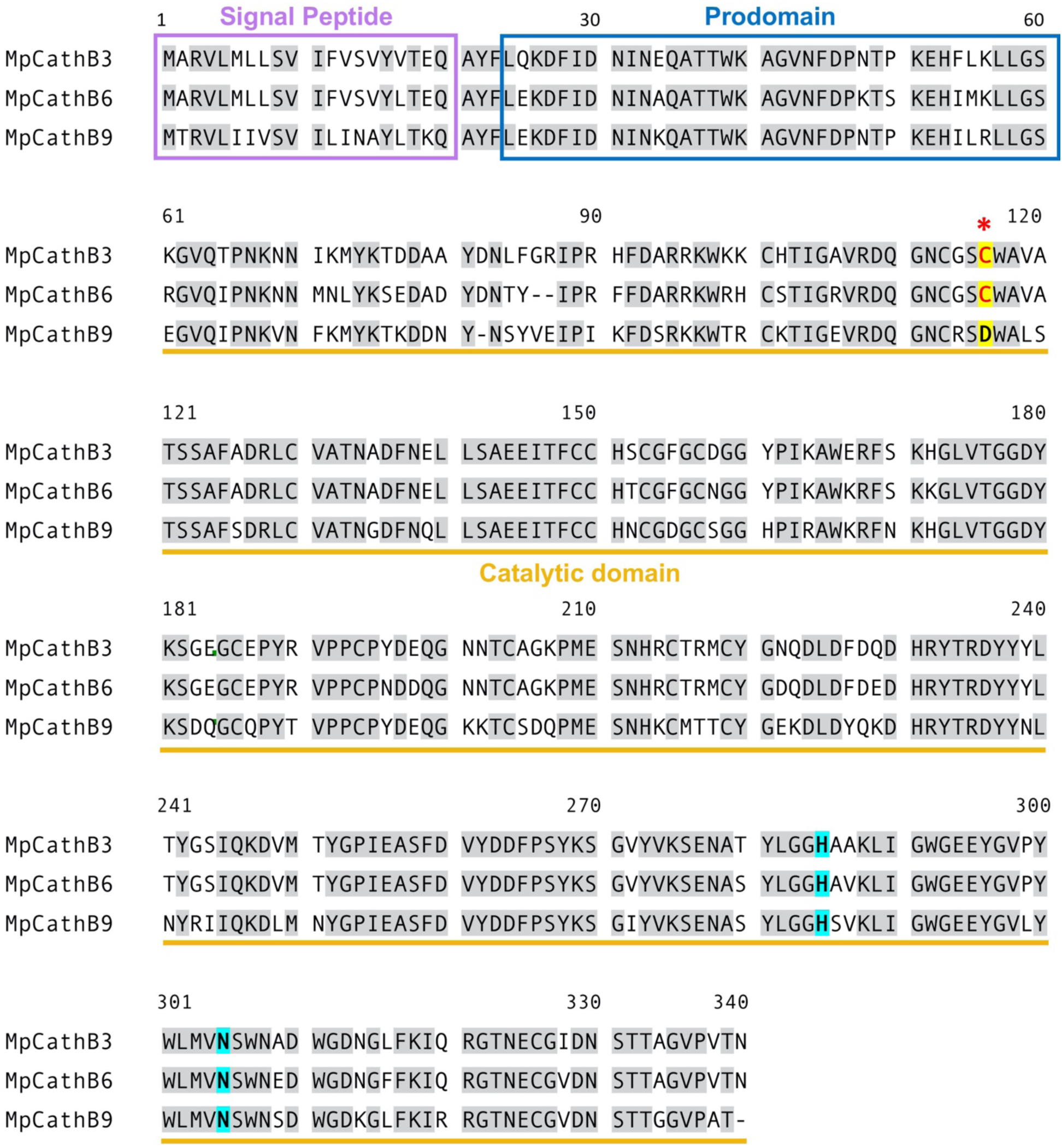
Multiple sequence alignment of *M. persicae* CathB3, 6 and 9 proteins using MUSCLE. The conserved residues are marked in grey background. Signal peptide and prodomain are labelled with rectangles. Catalytic domain is marked with a yellow line underneath the alignment. The catalytic cysteine is indicated as red font in yellow background and marked with a red asterisk. The catalytic histidine and asparagine are highlighted in cyan background.

**Fig. S5.**
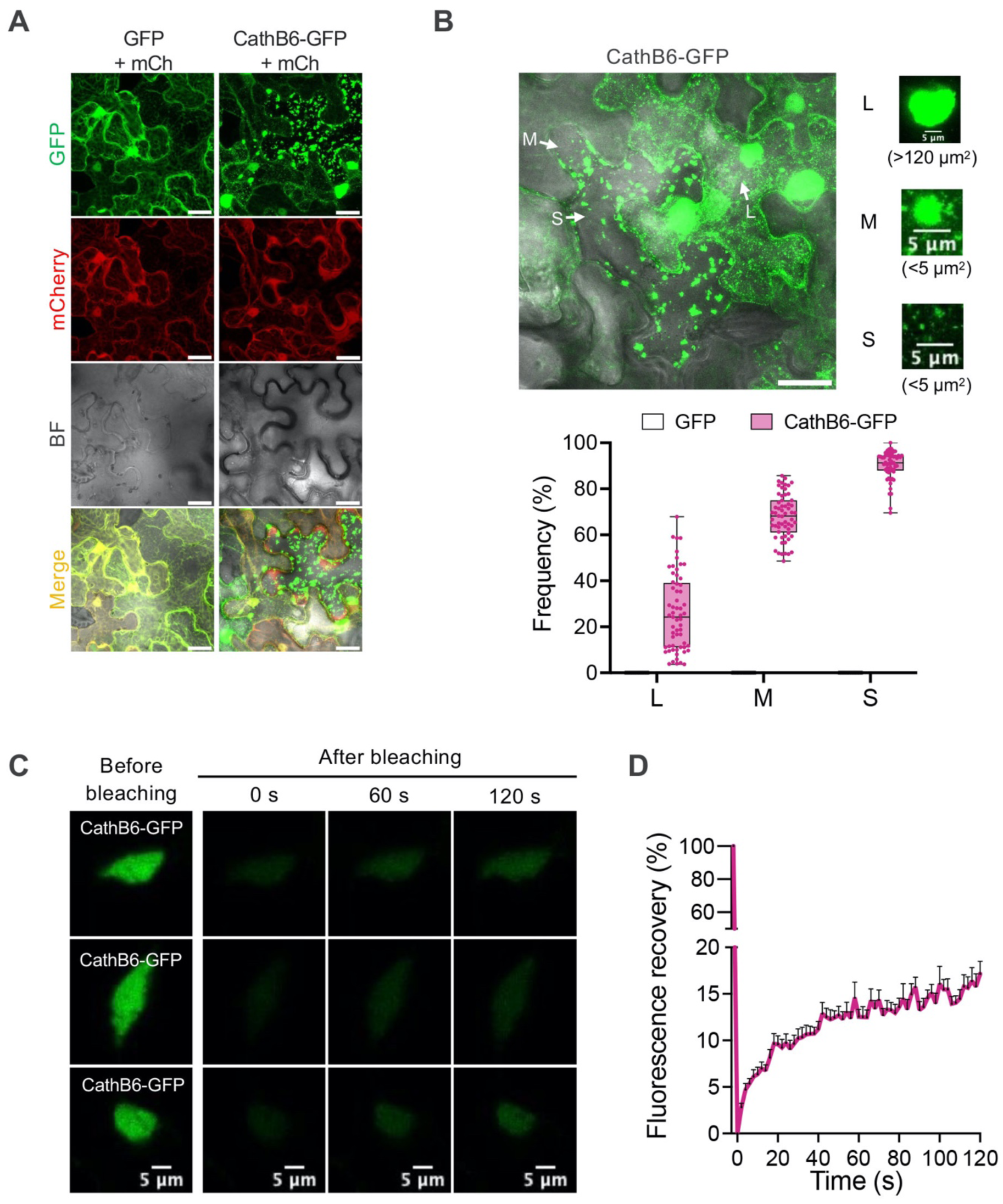
*M. persicae* CathB6 locates to mobile puncta within plant cells. (**A**) Confocal images of GFP and CathB6-GFP in *N. benthamiana* cells with mCherry serving as a reference marker. (**B**) Classification of CathB6-GFP puncta into three size categories: Large (L), Medium (M), and Small (S) based on area. Puncta frequency was determined as the ratio of cells with CathB6-GFP puncta to the total number of cells observed in the field. Data were compiled from 61 observation fields across three independent experiments, with 22–40 cells per field. Scale bars in (A) and (B), 30 μm. (**C**) Fluorescence recovery after photobleaching (FRAP) analysis of CathB6-GFP puncta. Representative FRAP images show fluorescence recovery over time, with Time 0 marking the photobleaching pulse. Data are representative of 25 independent experiments. (**D**) Time-course plot of fluorescence recovery for CathB6-GFP puncta after photobleaching. Recovery was quantified as the percentage of fluorescence intensity post-bleaching relative to pre-bleaching intensity. Data are presented as mean ± s.e.m. (n = 25).

**Fig. S6.**
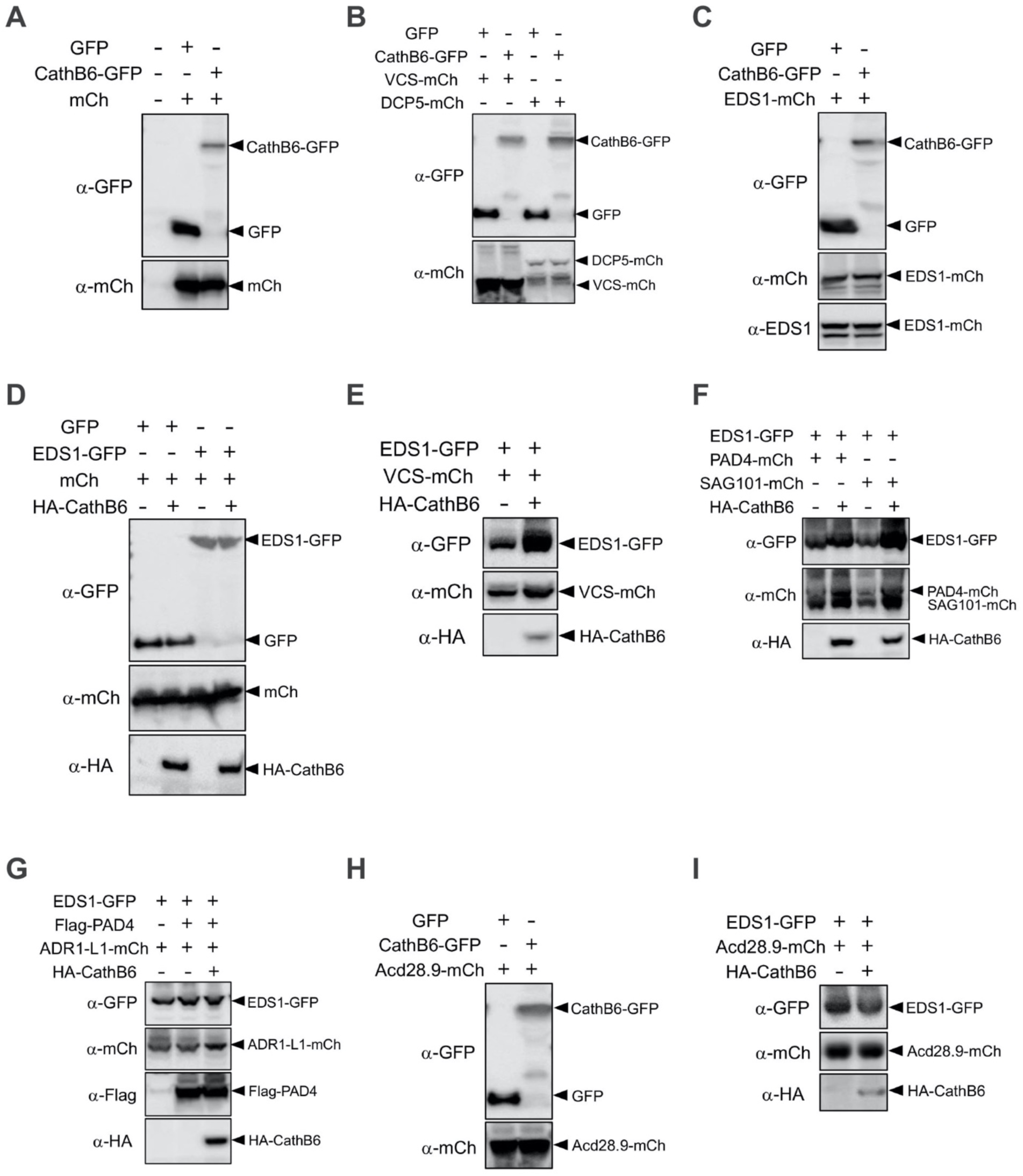
Western blots showing the presence of expected proteins in agroinfiltrated *N. benthamiana* leaf sections used for confocal microscopy experiments. (**A**) GFP, CathB6-GFP and mCherry, relates to Fig. 2A and fig. S5. (**B**) GFP, CathB6-GFP, VCS-mCherry and DCP5-mCherry, relates to Fig. 2, C and D, and fig. S8, A and B. (**C**) GFP, CathB6-GFP and EDS1-mCherry, relates to Fig. 3C. (**D**) GFP, EDS1-GFP, mCherry and HA-CathB6, relates to fig. S15A. (**E**) EDS1-GFP, VCS-mCherry and HA-CathB6, relates to Fig. 3D. (**F**) EDS1-GFP, PAD4-mCherry, SAG101-mCherry and HA-CathB6, relates to Fig. 3E and fig. S15B. (**G**) EDS1-GFP, Flag-PAD4, ADR1-L1-mCherry and HA-CathB6, relates to Fig. 3F. (**H**) GFP, CathB6-GFP and Acd28.9-mCherry, relates to Fig. 5B. (**I**) EDS1-GFP, Acd28.9-mCherry and HA-CathB6, relates to Fig. 5C.

**Fig. S7.**
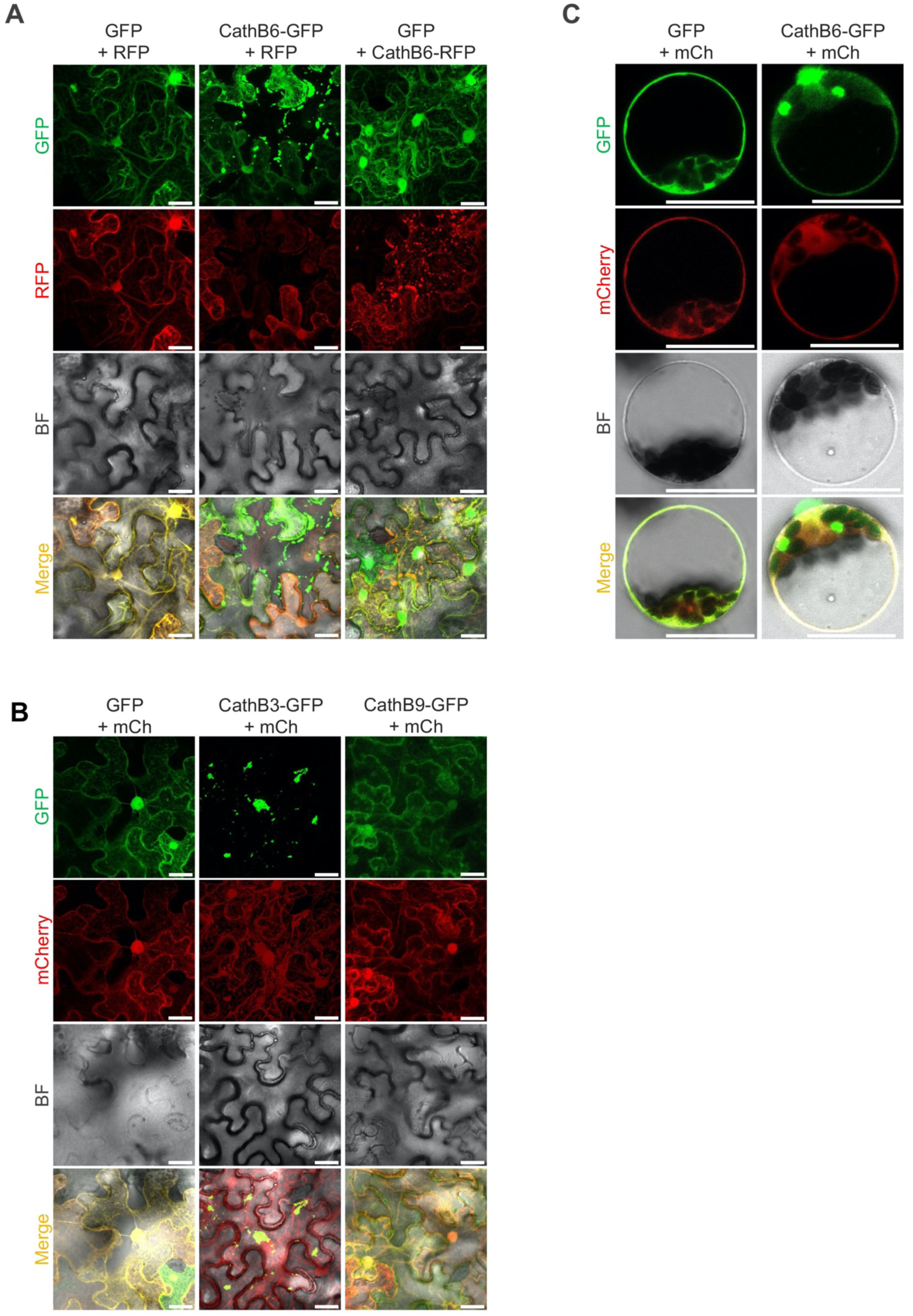
*M. persicae* CathB6 and CathB3, and to a lesser extent CathB9, form cytoplasmic puncta in *N. benthamiana* epidermal cells and *A. thaliana* protoplasts. (**A**) CathB6 forms puncta in *N. benthamiana* cells when fused to GFP (CathB6-GFP) or RFP (CathB6-RFP), with free RFP or GFP as internal references. (**B**) CathB3-GFP forms puncta of varying sizes in *N. benthamiana* cells, whereas CathB9-GFP generates significantly fewer puncta. (**C**) CathB6-GFP forms puncta of varying sizes in *A. thaliana* protoplasts. Scale bars, 30 μm.

**Fig. S8.**
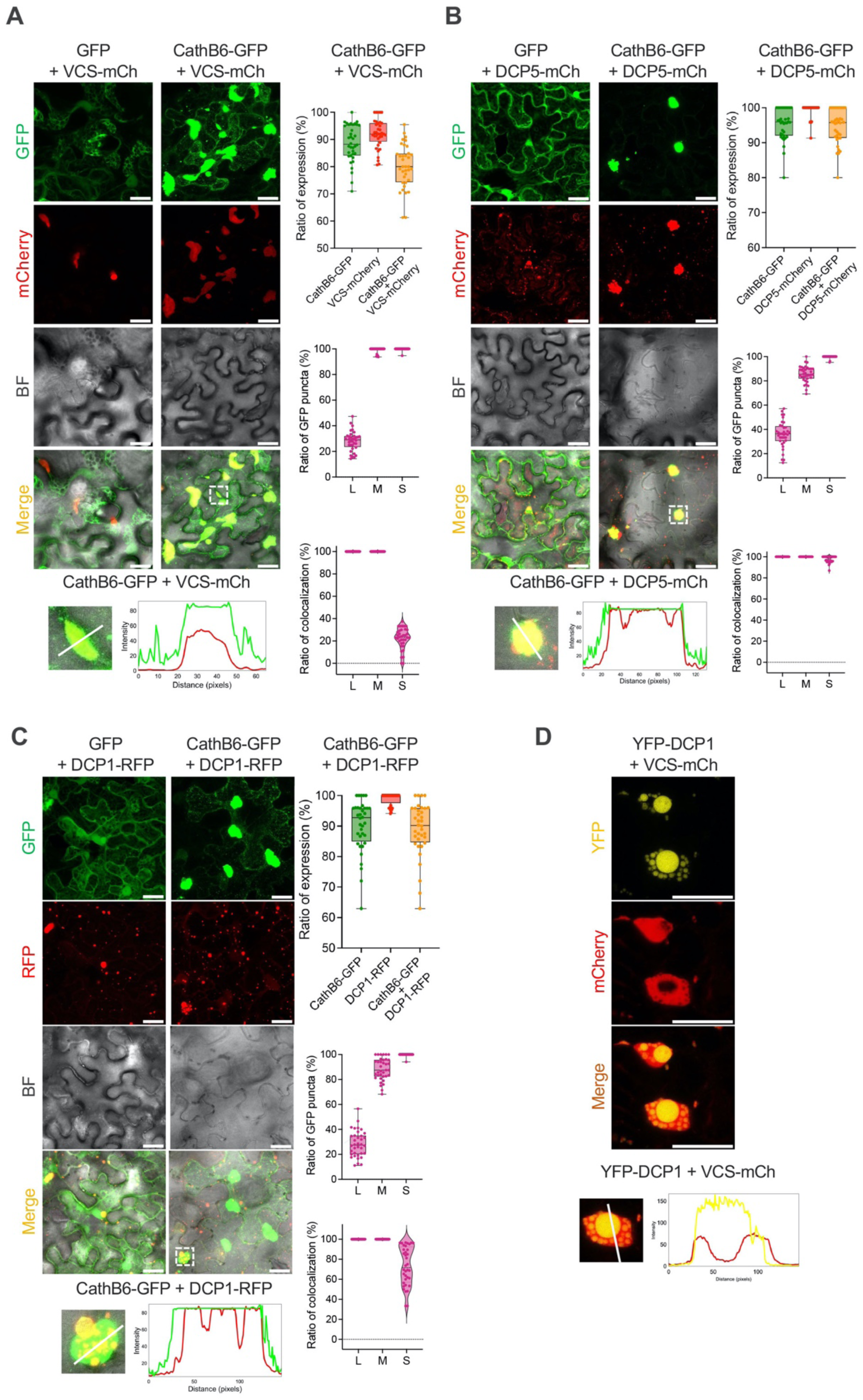
CathB6-GFP colocalizes with p-body markers VCS, DCP5 and DCP1 in plant cells. (**A** to **C**) CathB6-GFP colocalizes with VCS-mCherry (A), DCP5-mCherry (B) and DCP1-RFP (C) in *N. benthamiana* cells, unlike GFP alone (control). Panels at right of the confocal images provide quantification of colocalization from three independent experiments across 36 observation fields. Top: ratios of cells expressing CathB6-GFP and/or VCS-mCherry/DCP5-mCherry/DCP1-RFP. Middle: proportions of cells with large, medium, and small CathB6-GFP puncta colocalized with VCS-mCherry/DCP5-mCherry/DCP1-RFP. Bottom: Proportions of VCS-mCherry/DCP5-mCherry/DCP1-RFP puncta colocalized with CathB6-GFP in cells with CathB6-GFP. (**D**) Confocal images of YFP-DCP1 and VCS-mCherry in *N. benthamiana* cells, showing YFP-DCP1 localizing within VCS-mCherry puncta. Graphs below the confocal images in (A) to (D) are intensity profiles along the marked lines of one of the puncta in the confocal image above. Scale bars, 30 μm.

**Fig. S9.**
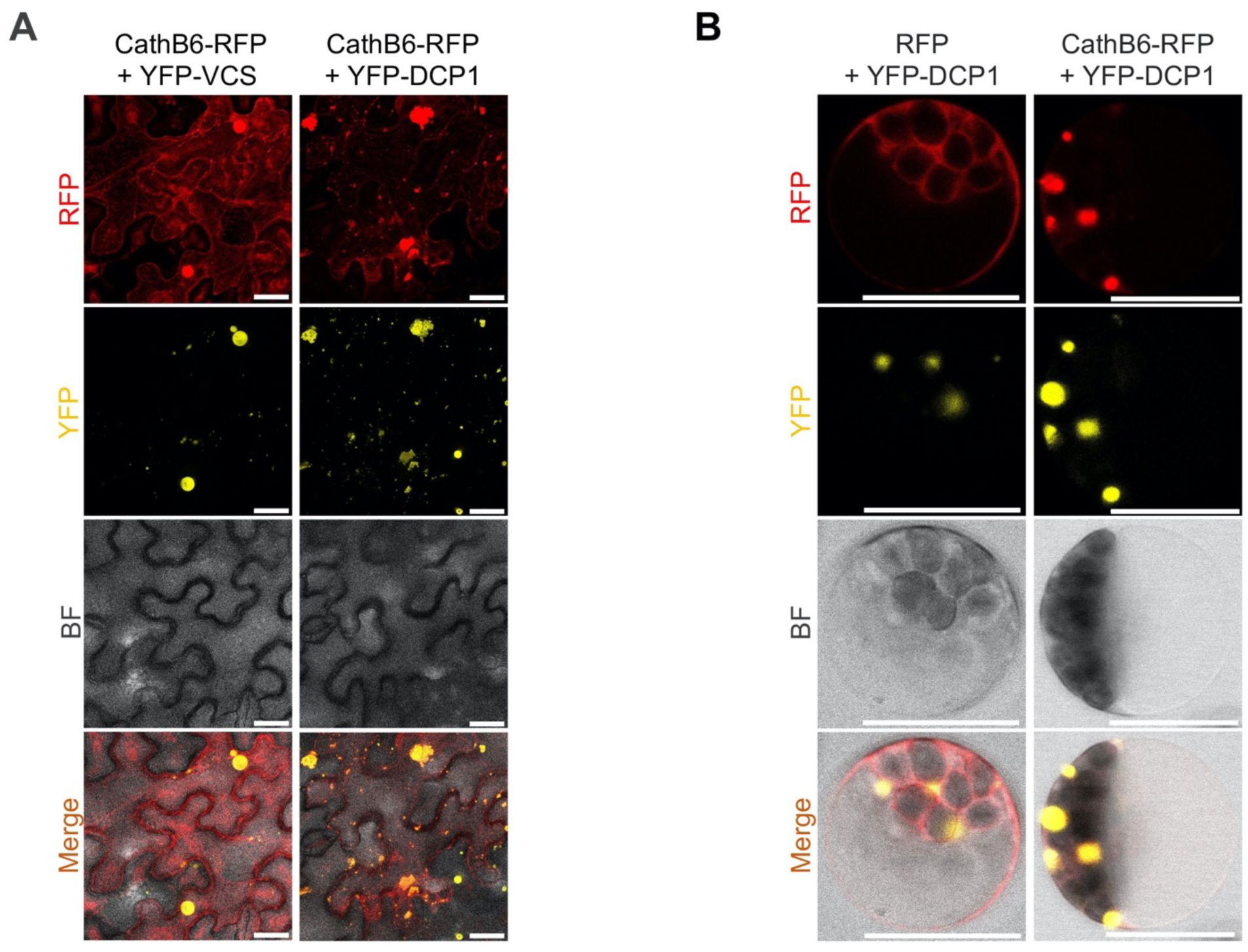
CathB6-RFP colocalizing with p-body markers YFP-VCS and YFP-DCP1 in *N. benthamiana* leaf cells (A) and *A. thaliana* protoplasts (B). Scale bars, 30 μm (A) and 20 μm (B).

**Fig. S10.**
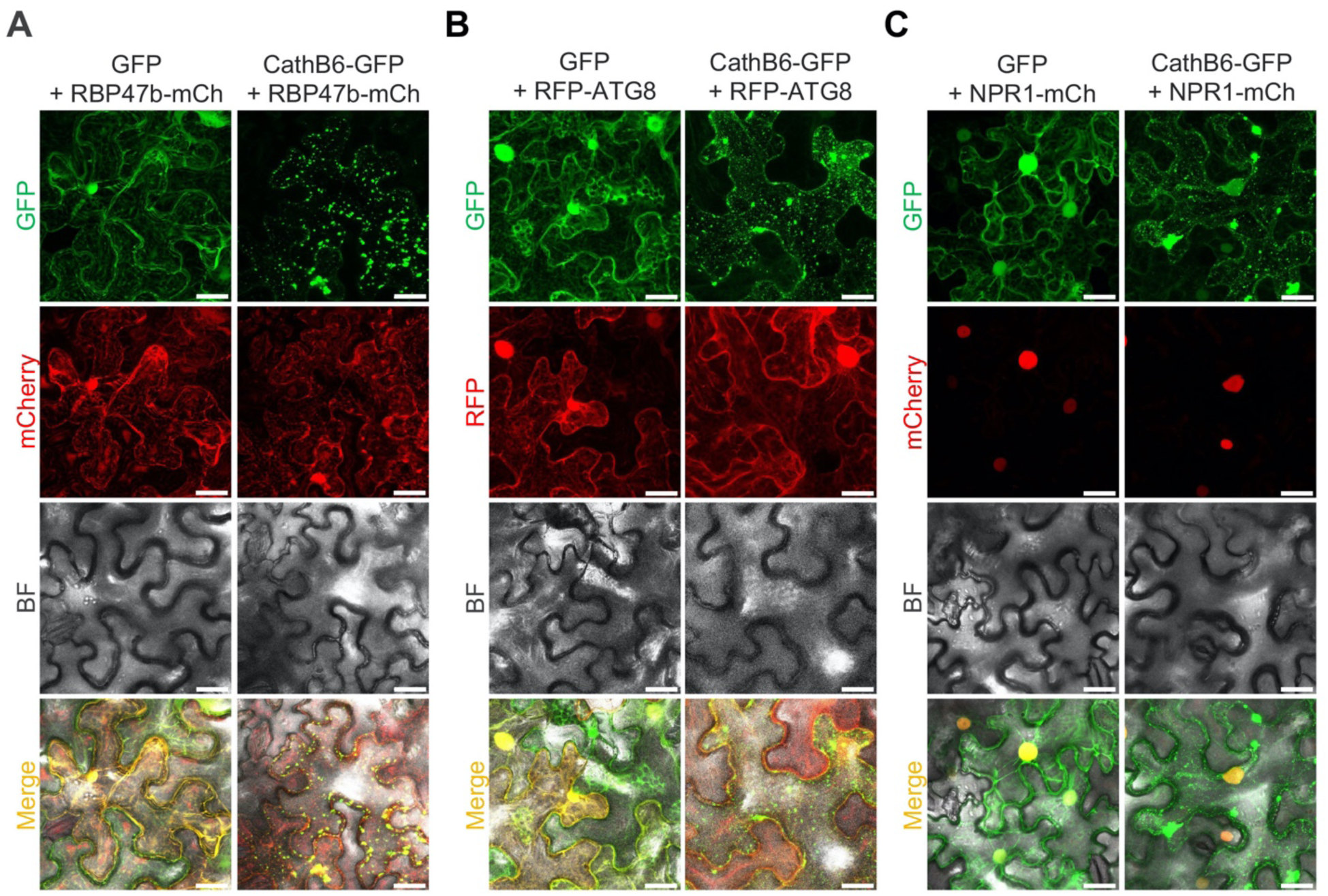
CathB6-GFP puncta partially colocalize with the stress granule marker RBP47b-mCherry (A), while no obvious colocalizations were observed with RFP-ATG8 (B) or NPR1-mCherry (C) in *N. benthamiana* epidermal cells. **Scale bars, 30 μm.**

**Fig. S11.**
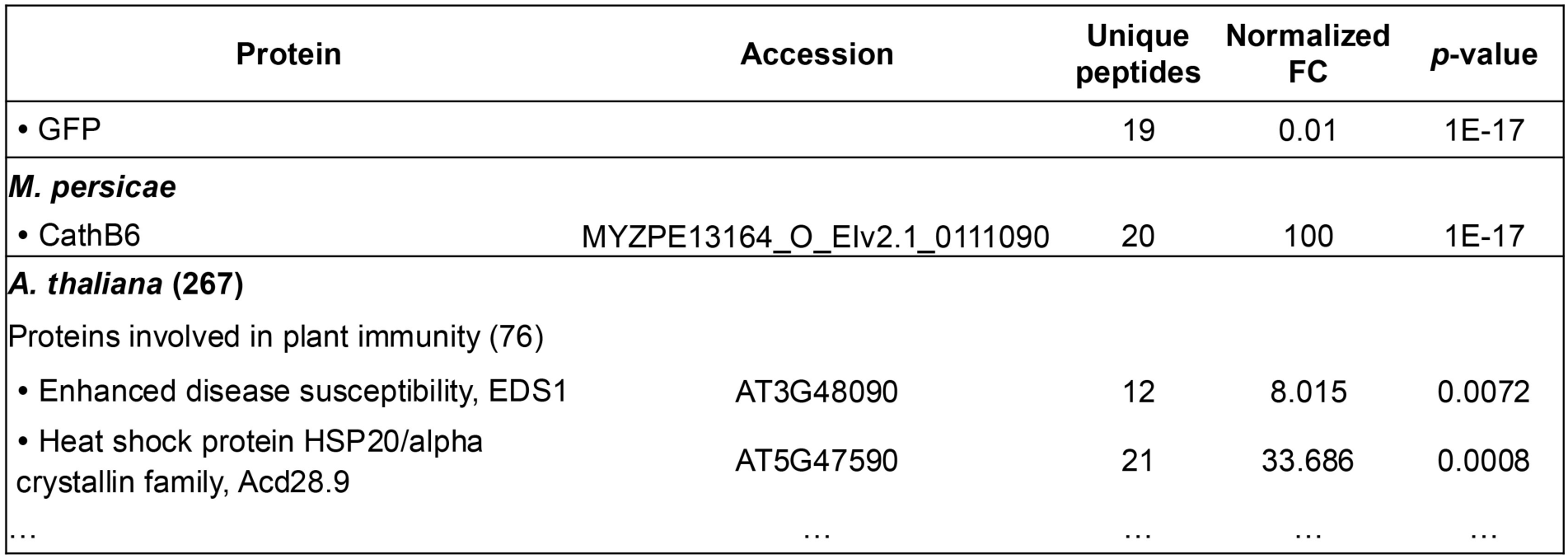
CathB6-TurboID PL-MS identifies EDS1 and Acd28.9 as CathB6 interactors in *A. thaliana*. *A. thaliana* proteins identified from CathB6-TurboID PL-MS were screened with a foldchange of CathB6-TurboID/GFP-TurboID > 2 and *p*-value < 0.05. Nineteen (19) unique peptides matching GFP were identified in the GFP-TurboID sample, and 20 matching CathB6 in the CathB6-GFP TurboID sample. Among a total of 267 *A. thaliana* proteins identified in the CathB6-GFP TurboID sample, 76 proteins are in involved in regulating plant immunity, and these include EDS1 and Acd28.9. Full CathB6-TurboID PL-MS dataset is shown in (Liu et al., 2024a).

**Fig. S12.**
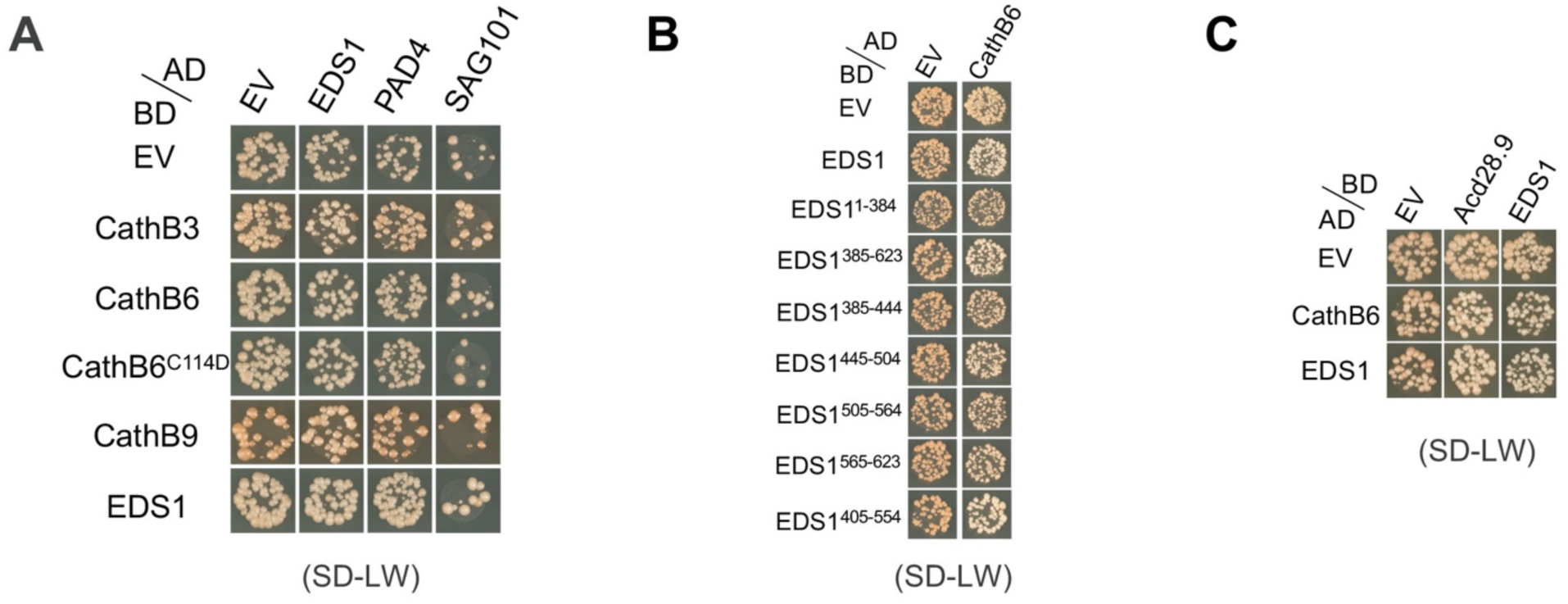
Yeast growth on double dropout medium lacking leucine and tryptophan indicating the expression of AD and BD constructs in Y2H assays. Related to Figure 3A (A), Figure 3B (B) and Figure 5A (C) of the main text.

**Fig. S13.**
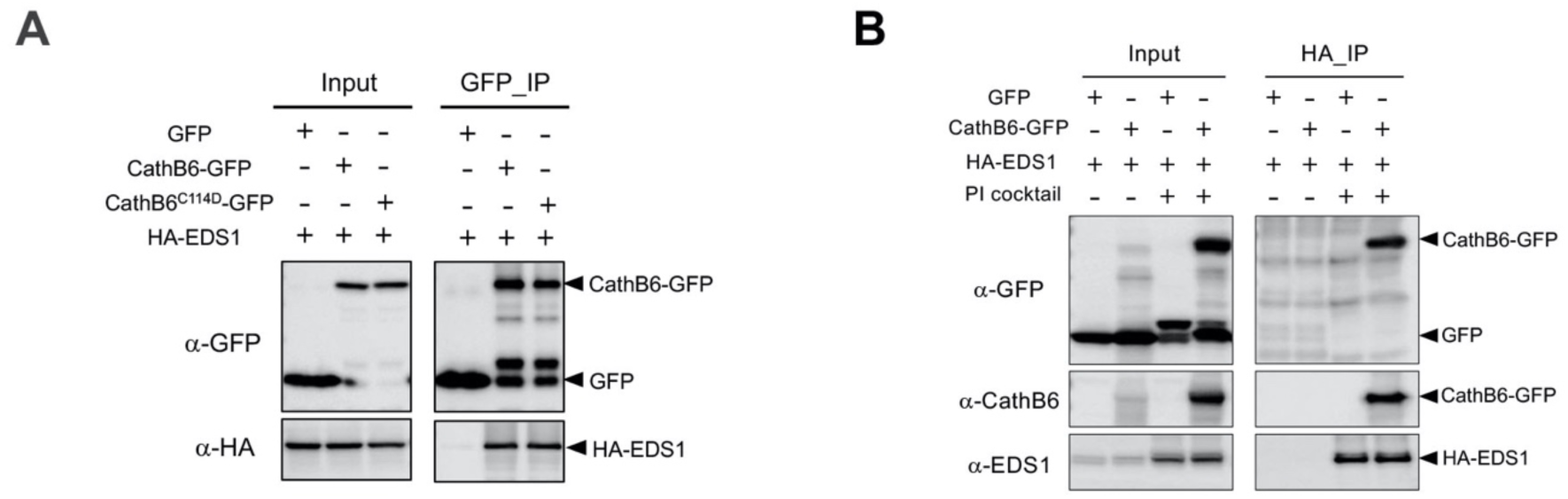
*M. persicae* CathB6 interacts with EDS1 *in planta* and *in vitro*. (**A**) Co-IP of HA-EDS1 with CathB6-GFP and CathB6^C114D^-GFP from agroinfiltrated *N. benthamiana* leaves using GFPtrap beads. (**B**) Co-IP of CathB6-GFP with HA-EDS1 from insect cell (Sf9) extracts, using anti-HA beads. The addition of protease inhibitor (PI) cocktail improved CathB6 and EDS1 stability in co-IP experiments. Arrowheads indicate bands of the expected sizes.

**Fig. S14.**
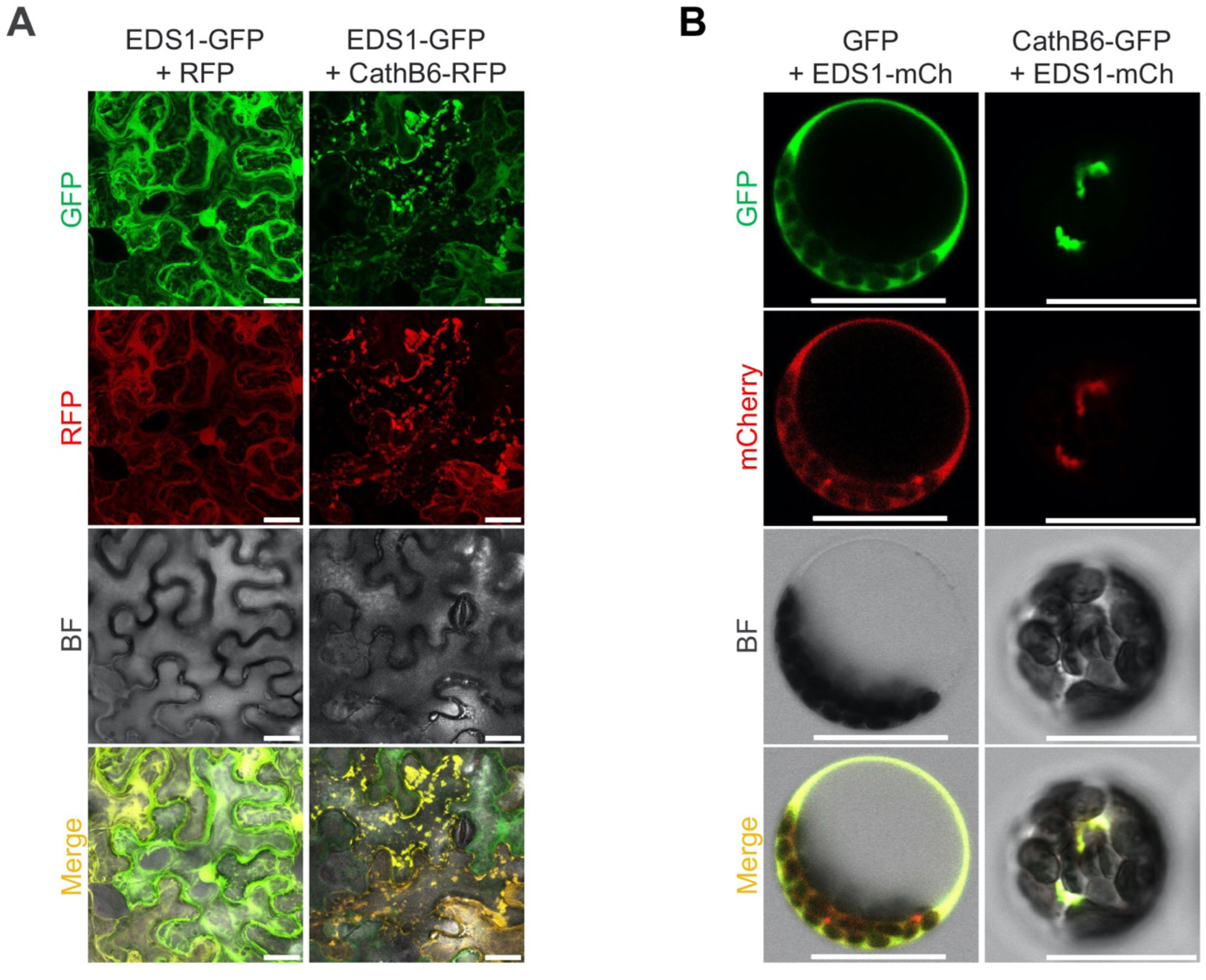
CathB6-RFP relocates EDS1-GFP to puncta in *N. benthamiana* epidermal cells (A) and *A. thaliana* protoplasts (B). Scale bars, 30 μm.

**Fig. S15.**
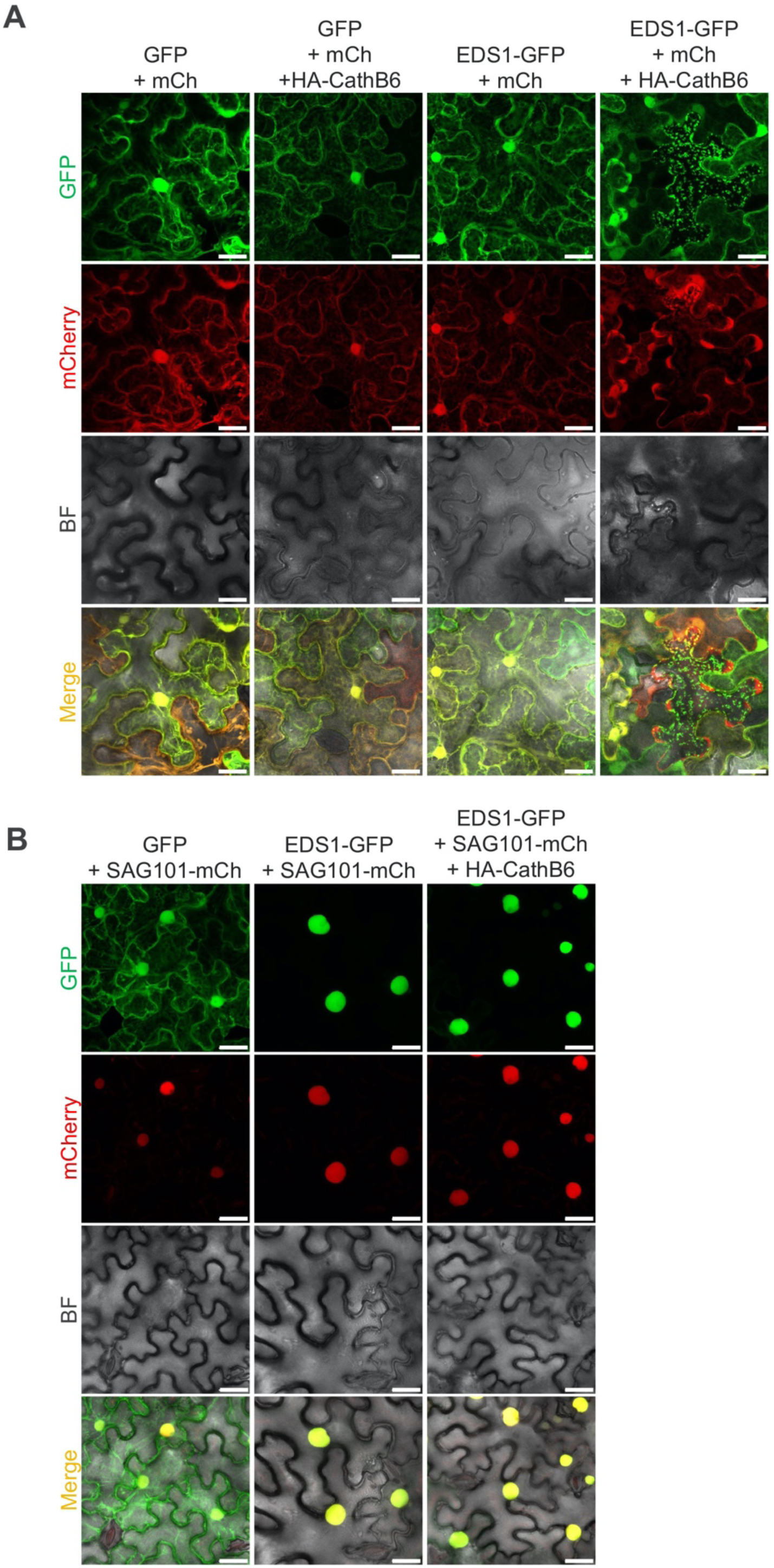
EDS1 alone localizes to cytoplasmic puncta/p-bodies (A), while the EDS1-SAG101 complex remains localized to nuclei (B), in the presence of *M. persicae* CathB6 in *N. benthamiana* leaves. Scale bars, 30 μm.

**Fig. S16.**
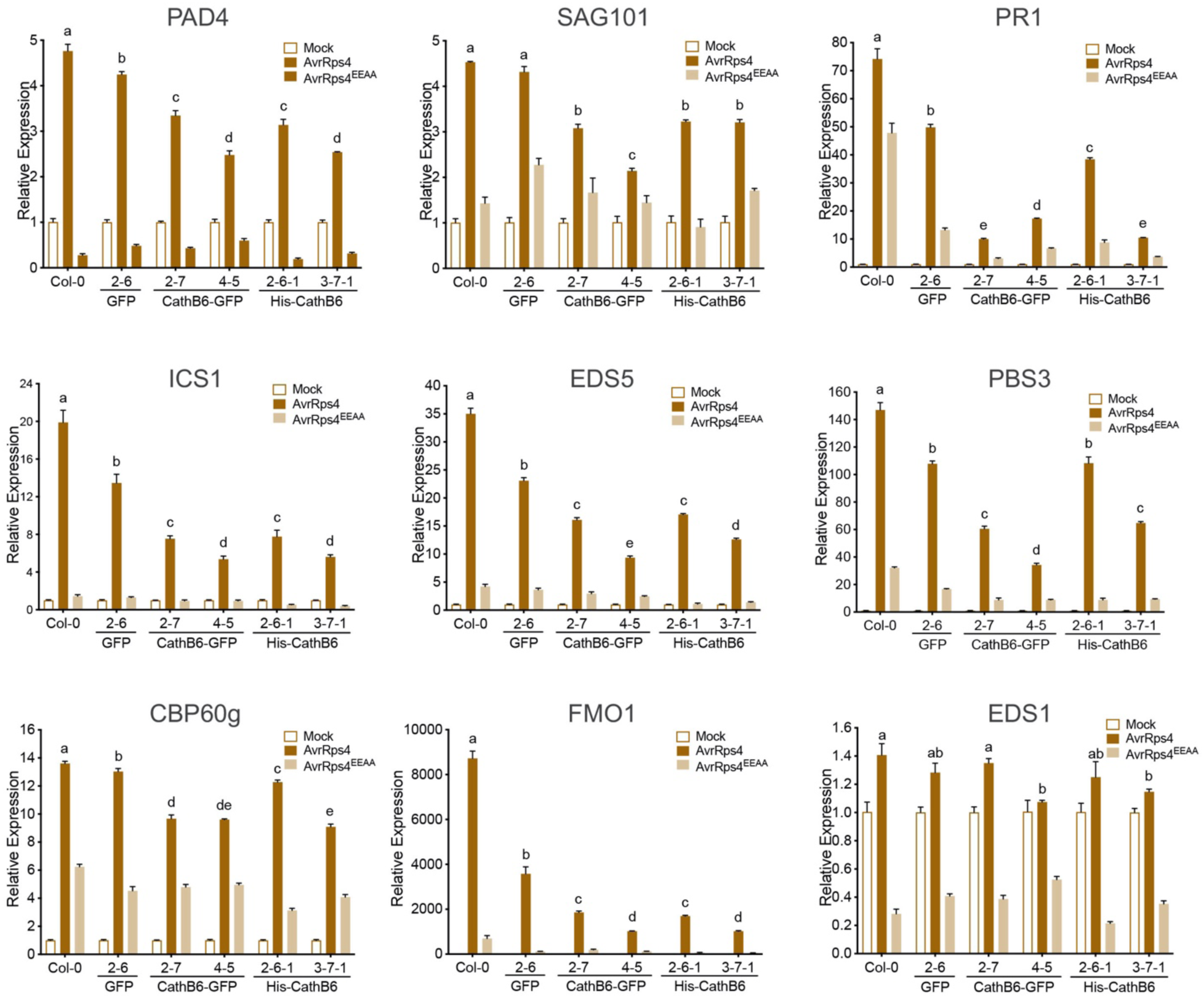
*M. persicae* CathB6 suppresses EDS1-responsive genes regulating systemic acquired resistance (SAR). Quantifications of transcript levels of EDS1 and SAR-related genes ICS1, EDS5, PBS3 (SA biosynthesis) and PAD4, SAG101, CBP60g, FMO1, PR1 (SAR regulators) in response to mock, Pf0-1 AvrRps4, or Pf0-1 AvrRps4^EEAA^ treatments using gene-specific primers. Data represent means ± s.e.m. from data of two biological replications; significant differences were analyzed by ANOVA. More replicates are shown in Fig. 4A.

**Fig. S17.**
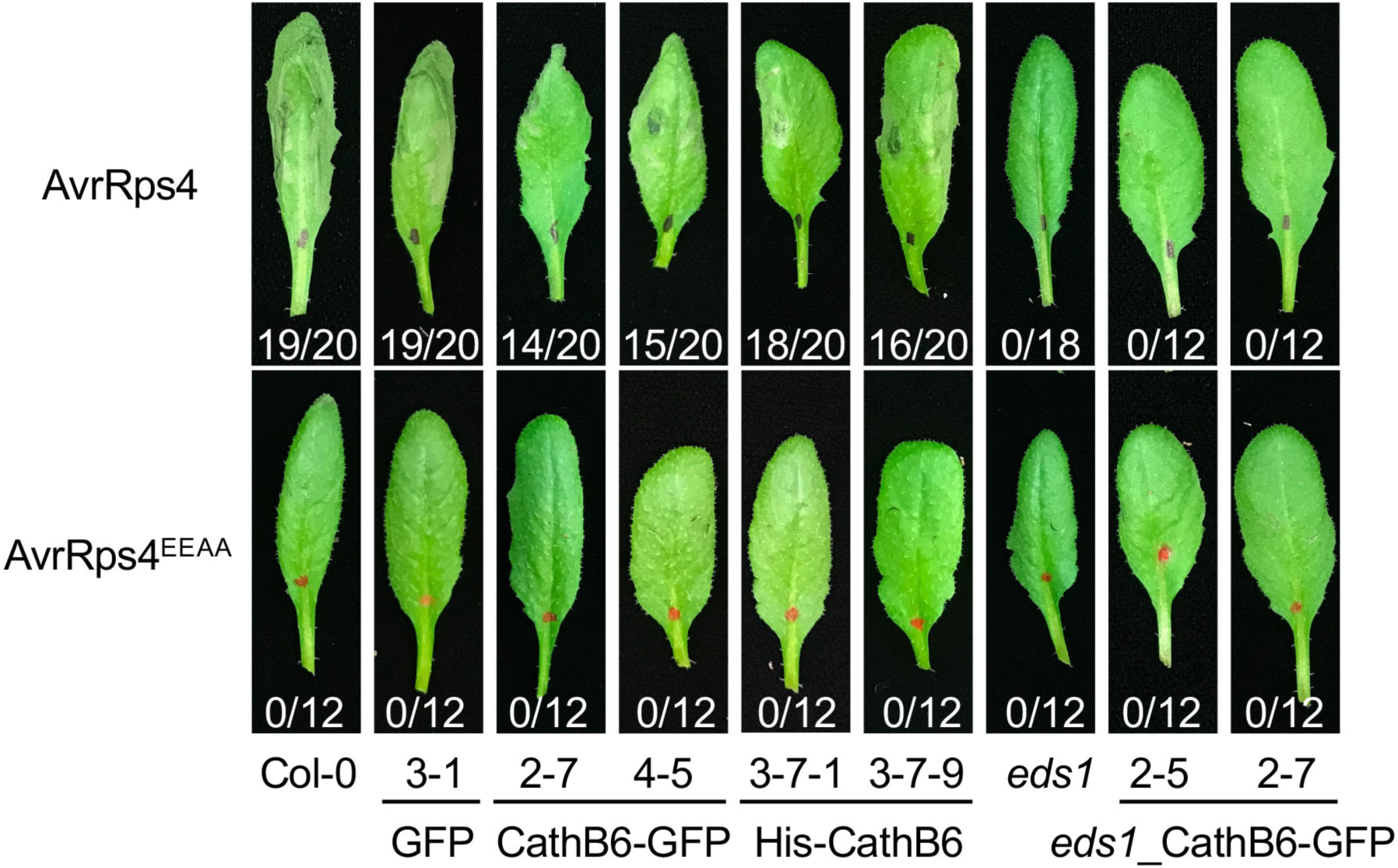
*M. persicae* CathB6 does not have obvious hypersensitive response (HR) suppression activity. Photos are representative leaves selected from 12 to 20 leaves of stable His-CathB6 or CathB6-GFP *A. thaliana* lines in Col-0 and *eds1-2* backgrounds, after infiltrating Pf0-1 AvrRps4 and AvrRps4^EEAA^ for 24 h. HR response typically seen upon infiltration with AvrRps4, but not AvrRps4^EEAA^, is evident from cell death symptoms on the leaves. The ratio beneath each leaf indicates number of leaves with visible tissue collapse in all infiltrated leaves from three independent experiments.

**Fig. S18.**
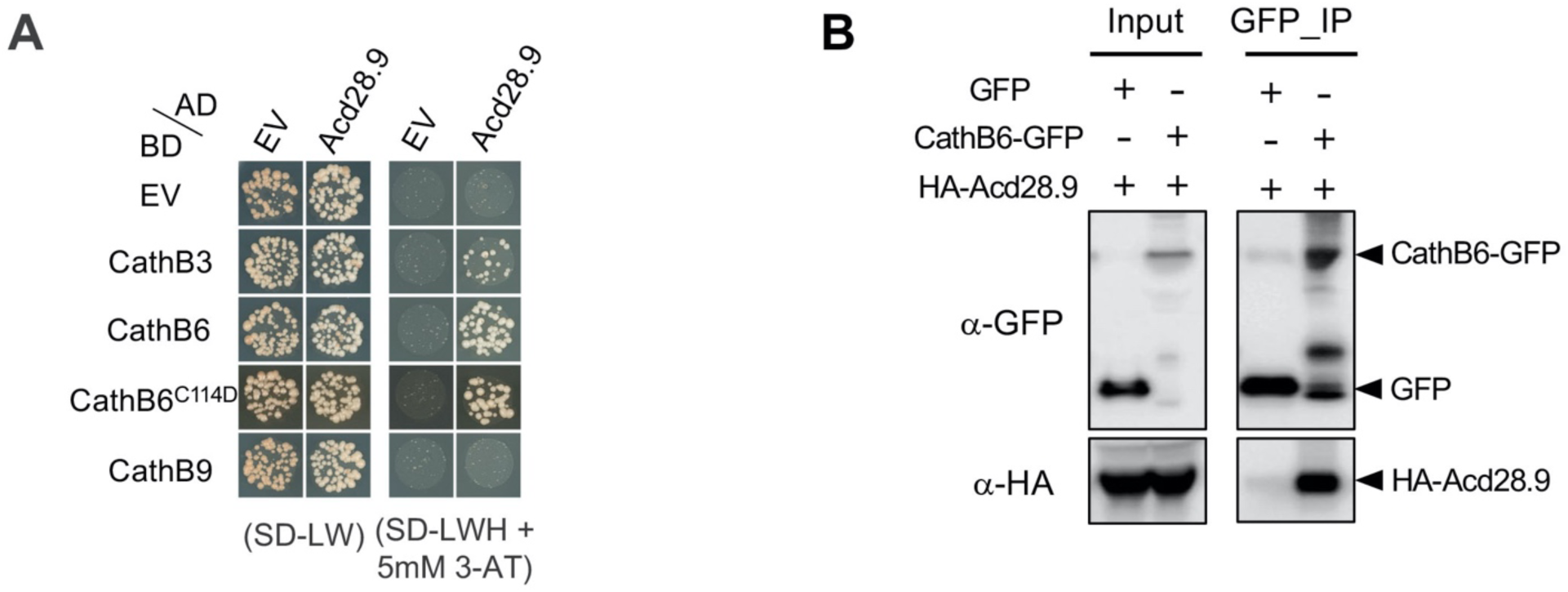
*M. persicae* CathB6 interacts with *A. thaliana* Hsp20 protein Acd28.9. (A) Yeast two-hybrid assays showing Acd28.9 interaction with CathB3, CathB6 and CathB6^C114D^. (**B**) Co-IP of HA-Acd28.9 with CathB6-GFP from agroinfiltrated *N. benthamiana* leaves using GFPtrap beads. Arrowheads indicate bands of the expected sizes.

**Fig. S19.**
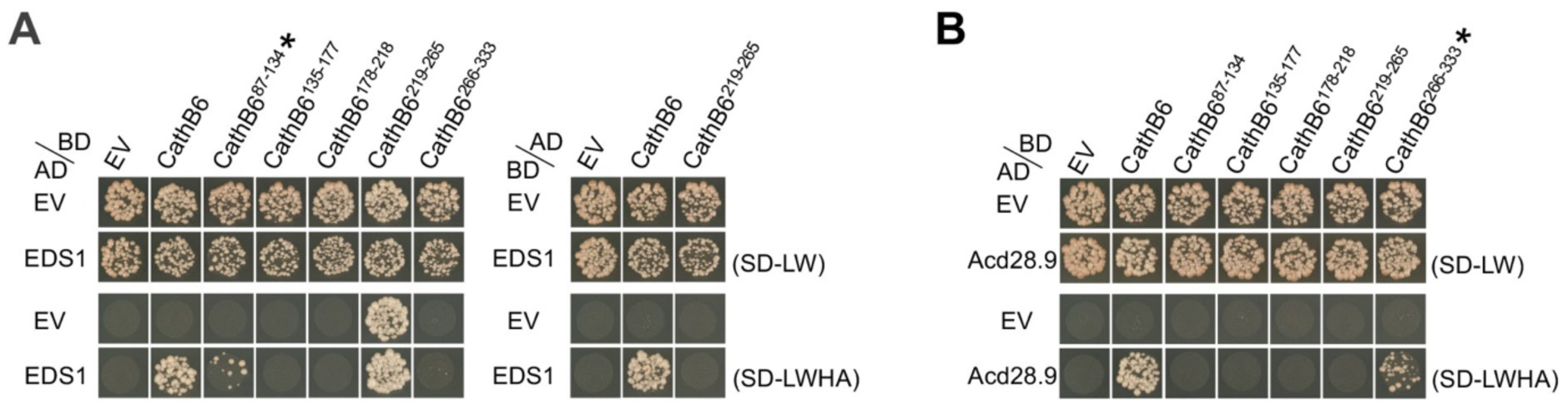
Yeast two hybrid assays showing that different fragments of *M. persicae* CathB6 interact with EDS1 (A) versus Acd28.9 (B). EV, empty vector control. AD, GAL4-activation domain. BD, GAL4-DNA binding domain. SD-LW, double dropout medium lacking leucine and tryptophan. SD-LWHA, quadruple dropout medium lacking leucine, tryptophan, histidine, and adenine. * indicates the fragment interacting with EDS1 (**A**) or Acd28.9 (**B**).

**Table S1.** List of CathB genes annotated in the v2 genome of the potato-peach aphid *Myzus persicae* clone O. DE, differentially expressed CathB genes in the *Myzus persicae* colonies reared on 8 divergent plant species (*Arabidopsis thaliana*, *Nicotiana benthamiana*, *Solanum tuberosum*, *Pisum sativum*, *Phaseolus vulgaris*, *Helianthus annuus*, *Chrysanthemum indicum* and *Zea mays*), compared to the ones reared on *Brassica rapa* (*13*).

| Name | Gene ID | Transcript ID | Expression on different plants | Note |
| --- | --- | --- | --- | --- |
| CathB1 | g2983.t1 | MYZPE13164_O_Elv2.1_0034760 | - |  |
| CathB2 | g2984.t2 | MYZPE13164_O_Elv2.1_0034770 | - |  |
| CathB3 | g7792.t1 | MYZPE13164_O_Elv2.1_0100520 | DE |  |
| CathB4 | g2985.t1 | MYZPE13164_O_Elv2.1_0034780 | - |  |
| CathB6 | g8486.t1 | MYZPE13164_O_Elv2.1_0111090 | DE |  |
| CathB7 | g6427.t1 | MYZPE13164_O_Elv2.1_0083010 | - |  |
| CathB8 | g8482.t1 | MYZPE13164_O_Elv2.1_0111040 | - |  |
| CathB9 | g8488.t1 | MYZPE13164_O_Elv2.1_0111110 | - |  |
| CathB10 | g8483.t1 | MYZPE13164_O_Elv2.1_0111050 | DE |  |
| CathB11 | g8484.t1 | MYZPE13164_O_Elv2.1_0111060 | - |  |
| CathB12 | g8485.t1 | MYZPE13164_O_Elv2.1_0111080 | DE |  |
| CathB13 | g8487.t1 | MYZPE13164_O_Elv2.1_0111100 | - |  |
| CathB14 | g8489.t1 | MYZPE13164_O_Elv2.1_0111120 | DE |  |
| CathB15 | g8490.t1 | MYZPE13164_O_Elv2.1_0111130 | DE |  |
| CathB16 | g27805.t1 | MYZPE13164_O_Elv2.1_0124680 | - | Truncated |
| CathB17 | g27806.t1 | MYZPE13164_O_Elv2.1_0124690 | DE |  |
| CathB18 | g27807.t1 | MYZPE13164_O_Elv2.1_0124700 | DE |  |
| CathB19 | g24530.t1 | MYZPE13164_O_Elv2.1_0159270 | DE |  |
| CathB20 | g24532.t1 | MYZPE13164_O_Elv2.1_0159280 | DE |  |
| CathB21 | g25994.t1 | MYZPE13164_O_Elv2.1_0179580 | - |  |
| CathB22 | g26516.t1 | MYZPE13164_O_Elv2.1_0186720 | - |  |
| CathB23 | g9913.t2 | MYZPE13164_O_Elv2.1_0269540 | DE |  |
| CathB24 | g10179.t1 | MYZPE13164_O_Elv2.1_0272820 | - |  |
| CathB25 | g12475.t1 | MYZPE13164_O_Elv2.1_0302340 | - |  |
| CathB26 | g12476.t1 | MYZPE13164_O_Elv2.1_0302350 | - |  |
| CathB27 | g12481.t1 | MYZPE13164_O_Elv2.1_0302390 | - |  |
| CathB28 | g12482.t1 | MYZPE13164_O_Elv2.1_0302400 | - |  |

**Table S2.**
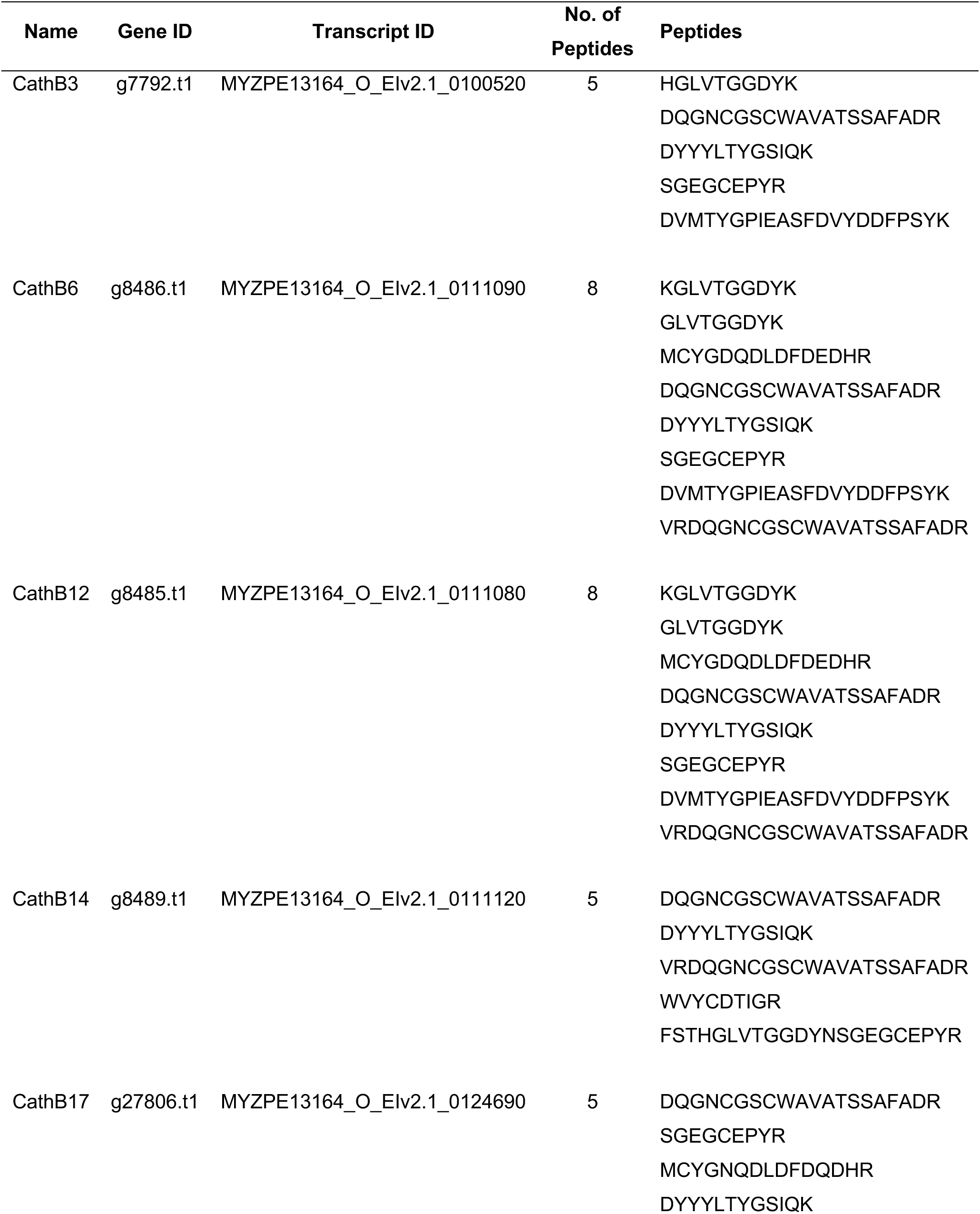
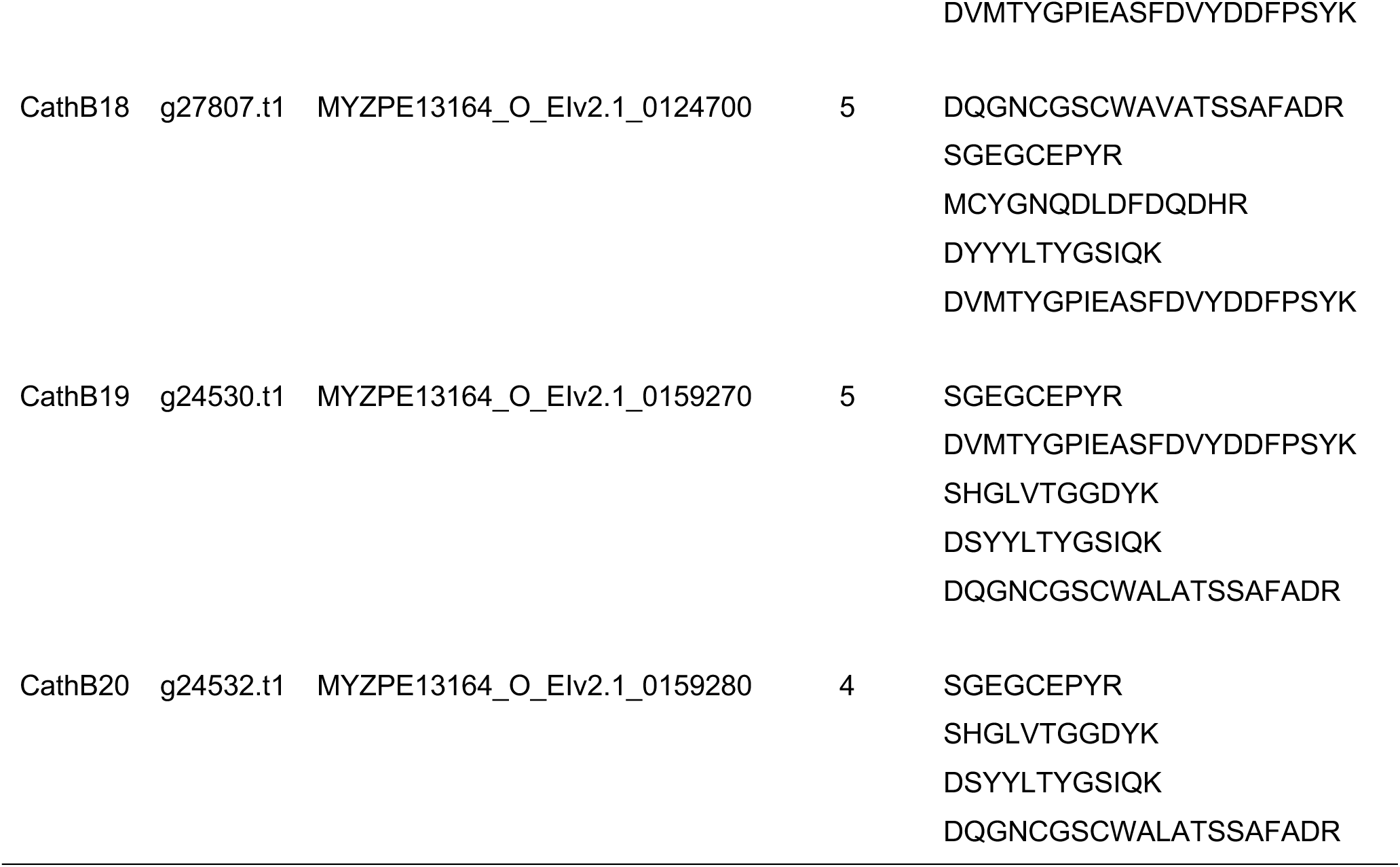
Peptides of CathB proteins detected in *M. persicae* Clone O oral secretion (OS) by MS. Gene ID and annotation ID of *M. persicae* clone O v2 database are labelled.

**Table S3.** Markers of cellular components used in the study.

| **Constructs** | **Protein name** | **Cellular components** | **References** |
| --- | --- | --- | --- |
| VCSc-mCherry | Varicose, VCS | Processing bodies | (Xu et al., 2006) |
| YFP-VCSc | Varicose, VCS | Processing bodies | (Petre et al., 2016) |
| DCP1-RFP | Decapping protein 1, DCP1 | Processing bodies | (Xu et al., 2006) |
| YFP-DCP1 | Decapping protein 1, DCP1 | Processing bodies | (Steffens et al., 2014) |
| DCP5-mCherry | Decapping protein 5, DCP5 | Processing bodies | (Xu and Chua, 2009) |
| RBP47b-mCherry | RNA binding protein 47b,<br>RBP47b | Stress granule | (Kosmacz et al., 2018) |
| RFP-ATG8 | Autophagy-related protein 8,<br>ATG8 | Autophagosome | (Pu and Bassham, 2023) |
| NPR1-mCherry | Nonexpressor of pathogenesis<br>related 1, NPR1 | NPR1 condensates | (Zavaliev et al., 2020) |

**Table S4.**
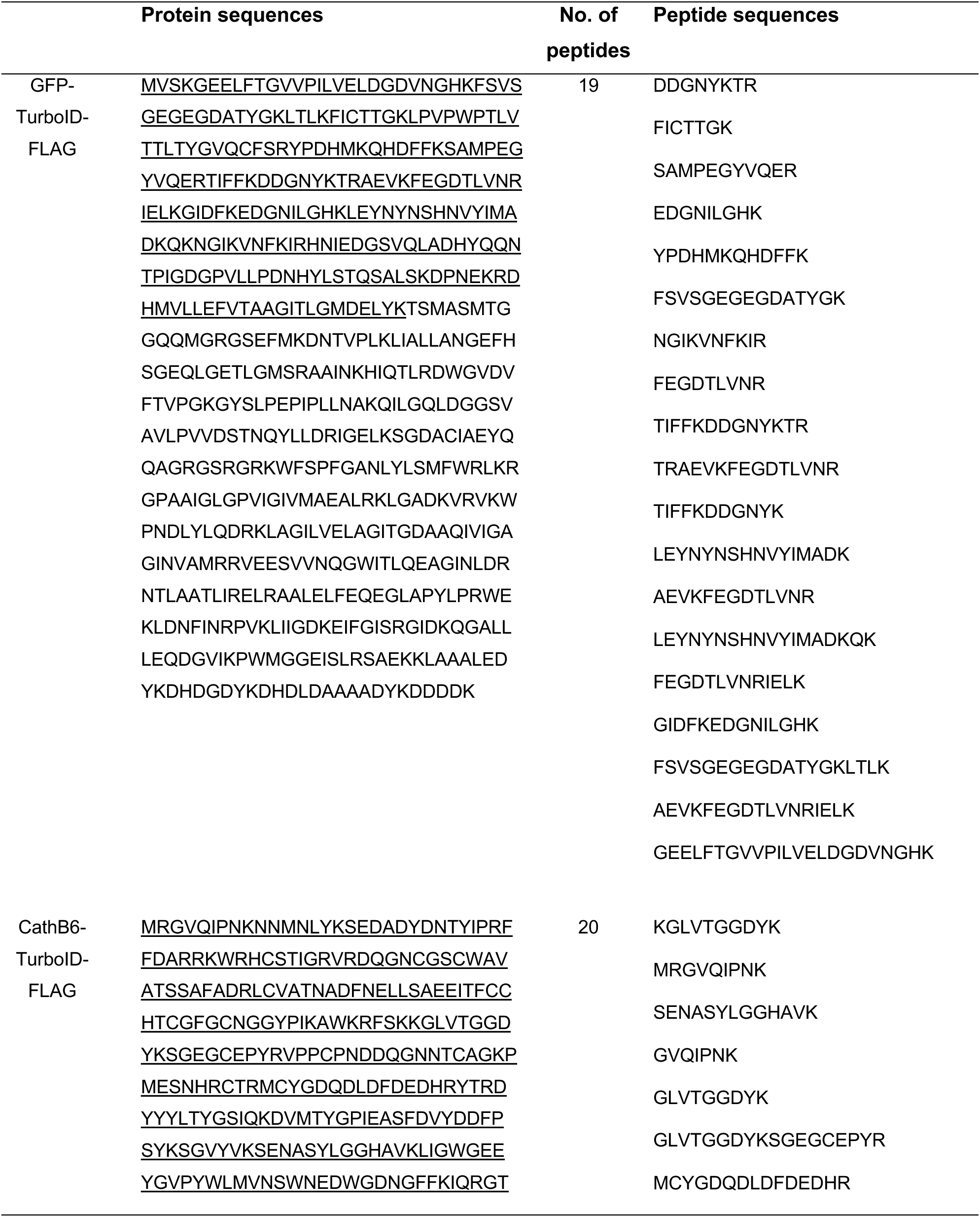
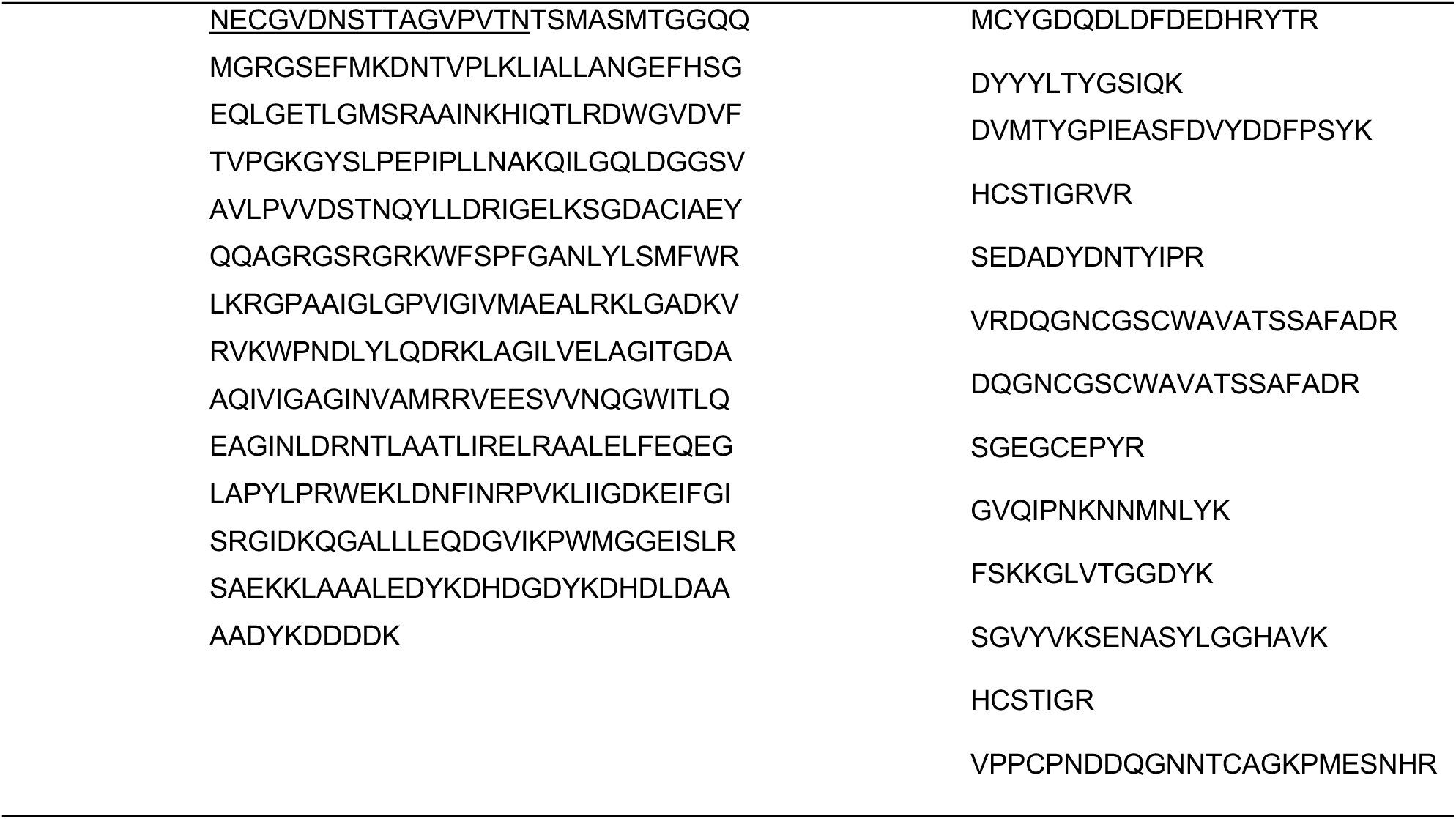
Peptides corresponding to GFP (GFP-TurboID-FLAG) or CathB6 (GFP-TurboID-FLAG) detected in the TurboID-based proximity samples by mass spectrometry. Amino acids corresponding to GFP and CathB6 sequences within the GFP-TurboID-FLAG and CathB6-TurboID-FLAG sequences are underlined.

**Table S5.**
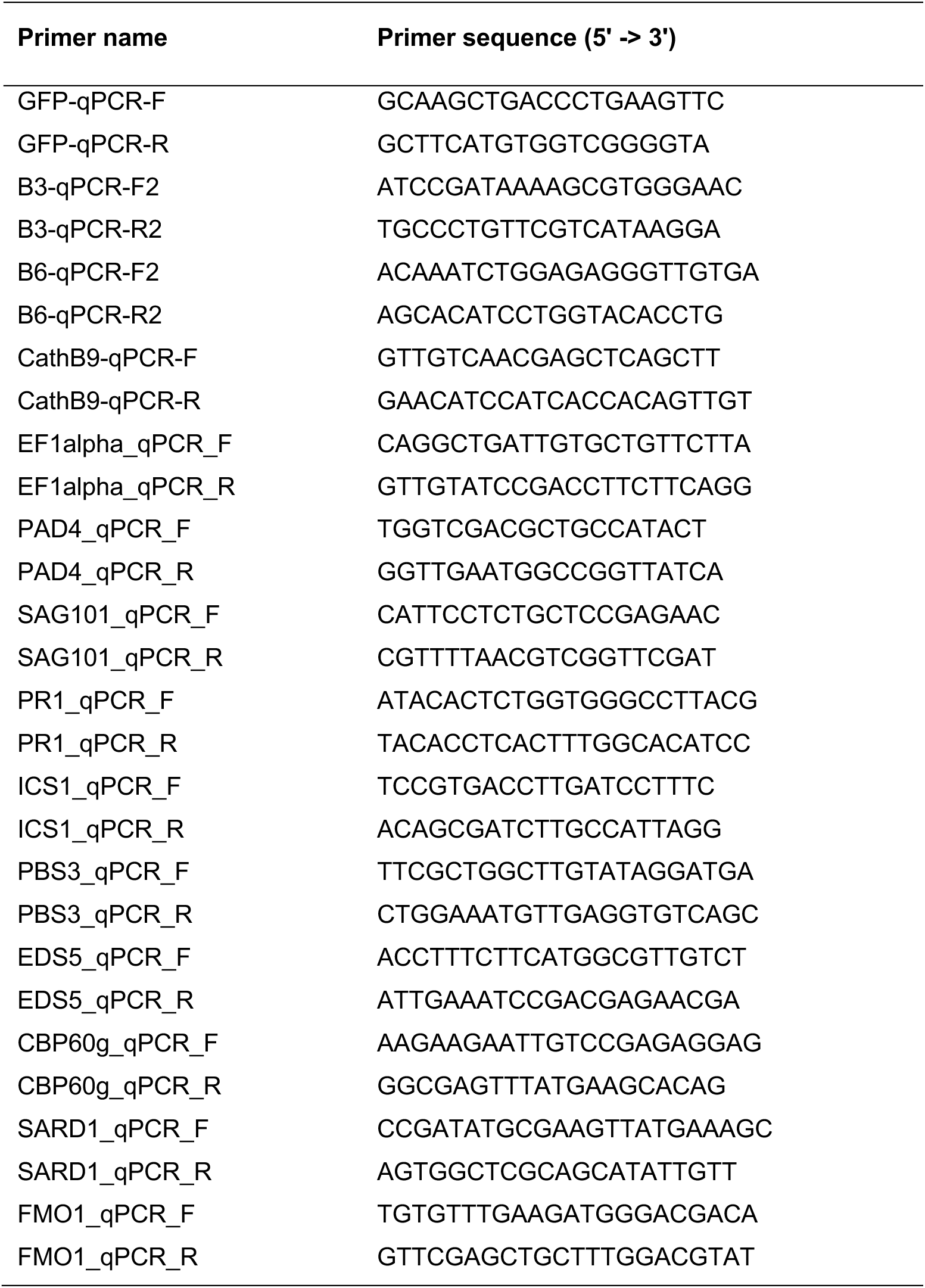
qRT-PCR primers used in the study.

**Table S6.** Plasmids used in the study.

| Plasmid name | Purpose | Ref. |
| --- | --- | --- |
| pBI121-CathB3 | Generating stable transgenic *A. thaliana* plants of His-CathB3 | This study |
| pBI121-CathB3<sup>C116D</sup> | Generating stable transgenic *A. thaliana* plants of His-CathB3<sup>C116D</sup> | This study |
| pBI121-CathB6 | Generating stable transgenic *A. thaliana* plants of His-CathB6 | This study |
| pBI121-CathB6<sup>C114D</sup> | Generating stable transgenic *A. thaliana* plants of His-CathB6<sup>C114D</sup> | This study |
| pBI121-CathB9 | Generating stable transgenic *A. thaliana* plants of His-CathB9 | This study |
| pB7FWG2-ATG | Generating stable transgenic *A. thaliana* plants of GFP, Transient expression of GFP in *A. thaliana* protoplast and *N. benthamiana* | This study |
| pB7RWG2-ATG | Transient expression of RFP in *A. thaliana* protoplast and *N. benthamiana* | This study |
| pB7WG2-mCherry | Transient expression of mCherry in *N. benthamiana* | This study |
| pB7FWG2-CathB3 | Transient expression of CathB3-GFP in *N. benthamiana* | This study |
| pB7FWG2-CathB6 | Generating stable transgenic *A. thaliana* plants of CathB6-GFP, Transient expression of CathB6-GFP in *A. thaliana* protoplast and *N. benthamiana* | This study |
| pB7FWG2-CathB6<sup>C114D</sup> | Transient expression of CathB6<sup>C114D</sup>-GFP in *N. benthamiana* | This study |
| pB7RWG2-CathB6 | Transient expression of CathB6-RFP in *A. thaliana* protoplast and *N. benthamiana* | This study |
| pB7FWG2-CathB9 | Generating stable transgenic *A. thaliana* plants of CathB9-GFP, Transient expression of CathB9-GFP in *N. benthamiana* | This study |
| pB7WG2-VCS-mCherry | Transient expression of VCS-mCherry in *N. benthamiana* | This study |
| pH7YWG2-YFP-VCS<sup>1</sup> | Transient expression of YFP-VCS in *N. benthamiana* | (Petre et al., 2016) |
| pB7RWG2-DCP1<sup>2</sup> | Transient expression of DCP1-RFP in *N. benthamiana* | (May et al., 2020) |
| pABM043-YFP-DCP1<sup>2</sup> | Transient expression of YFP-DCP1 in *A. thaliana* protoplast and *N. benthamiana* | (Merret et al., 2013) |
| pB7WG2-DCP5-mCherry | Transient expression in *N. benthamiana* | This study |
| pB7WG2-Acd28.9-mCherry | Transient expression of Acd28.9-mCherry in *N. benthamiana* | This study |
| pB7WG2-GFP-TurboID | Generating stable transgenic plants of GFP-TurboID-FLAG | This study |
| pB7WG2-CathB6-TurboID | Generating stable transgenic plants of CathB6-TurboID-FLAG | This study |
| pB7WG2-EDS1-mCherry | Transient expression of EDS1-mCherry in *A. thaliana* protoplast and *N. benthamiana* | This study |
| pB7WG2-PAD4-mCherry | Transient expression of PAD4-mCherry in *N. benthamiana* | This study |
| pB7WG2-SAG101-mCherry | Transient expression of SAG101-mCherry in *N. benthamiana* | This study |
| pB7WG2-ADR1-L1-mCherry<sup>3</sup> | Transient expression of ADR1-L1-mCherry in *N. benthamiana* | (Wang et al., 2023) |
| pB7FWG2-EDS1 | Transient expression of EDS1-GFP in *N. benthamiana* | This study |
| pB7WG2-RBP47b-mCherry | Transient expression of RBP47b-mCherry in *N. benthamiana* | This study |
| pB7WG2-RFP-ATG8 | Transient expression of RFP-ATG8 in *N. benthamiana* | (Leong et al., 2022) |
| pB7WG2-NPR1-mCherry | Transient expression of NPR1-mCherry in *N. benthamiana* | This study |
| pB7WG2-HA-CathB6 | Transient expression of HA-CathB6 in *N. benthamiana* | This study |
| pB7WG2-HA-Acd28.9 | Transient expression of HA-Acd28.9 in *N. benthamiana* | This study |
| pB7WG2-HA-EDS1 | Transient expression of HA-EDS1 in *N. benthamiana* | This study |
| pB7WG2-Flag-PAD4 | Transient expression of Flag-PAD4 in *N. benthamiana* | This study |
| pDESTGBKT7 | Y2H Empty vector of BD | This study |
| pDESTGADT7 | Y2H Empty vector of AD | This study |
| pDESTGBKT7-CathB3 | Y2H analysis of BD-CathB3 | This study |
| pDESTGBKT7-CathB6 | Y2H analysis of BD-CathB6 | This study |
| pDESTGBKT7-CathB6<sup>C114D</sup> | Y2H analysis of BD-CathB6<sup>C114D</sup> | This study |
| pDESTGBKT7-CathB6<sup>C114A</sup> | Y2H analysis of BD-CathB6<sup>C114A</sup> | This study |
| pDESTGBKT7-CathB9 | Y2H analysis of BD-CathB9 | This study |
| pDESTGBKT7-AtCathB3 | Y2H analysis of BD-AtCathB3 | This study |
| pDESTGBKT7-AtCathB3<sup>C113A</sup> | Y2H analysis of BD-AtCathB3<sup>C113A</sup> | This study |
| pDESTGBKT7-EDS1 | Y2H analysis of BD-EDS1 | This study |
| pDESTGBKT7-Acd28.9 | Y2H analysis of BD-Acd28.9 | This study |
| pDESTGADT7-CathB6 | Y2H analysis of AD-CathB6 | This study |
| pDESTGADT7-CathB6<sup>C114A</sup> | Y2H analysis of AD-CathB6<sup>C114A</sup> | This study |
| pDESTGADT7-AtCathB3 | Y2H analysis of AD-AtCathB3 | This study |
| pDESTGADT7-AtCathB3<sup>C113A</sup> | Y2H analysis of AD-AtCathB3<sup>C113A</sup> | This study |
| pDESTGADT7-EDS1 | Y2H analysis of AD-EDS1 | This study |
| pDESTGADT7-EDS1<sup>1-384</sup> | Y2H analysis of AD-EDS1<sup>1-384</sup> | This study |
| pDESTGADT7-EDS1<sup>385-623</sup> | Y2H analysis of AD-EDS1<sup>385-623</sup> | This study |
| pDESTGADT7-EDS1<sup>385-444</sup> | Y2H analysis of AD-EDS1<sup>385-444</sup> | This study |
| pDESTGADT7-EDS1<sup>445-504</sup> | Y2H analysis of AD-EDS1<sup>445-504</sup> | This study |
| pDESTGADT7-EDS1<sup>505-564</sup> | Y2H analysis of AD-EDS1<sup>505-564</sup> | This study |
| pDESTGADT7-EDS1<sup>565-623</sup> | Y2H analysis of AD-EDS1<sup>565-623</sup> | This study |
| pDESTGADT7-EDS1<sup>405-554</sup> | Y2H analysis of AD-EDS1<sup>405-554</sup> | This study |
| pDESTGADT7-PAD4 | Y2H analysis of AD-PAD4 | This study |
| pDESTGADT7-SAG101 | Y2H analysis of AD-SAG101 | This study |
| pFastBac-HTB-GFP | Expression of GFP in Sf9 cells using baculovirus system | This study |
| pFastBac-HTB-CathB6-GFP | Expression of CathB6-GFP in Sf9 cells using baculovirus system | This study |
| pFastBac-HTB-HA-EDS1 | Expression of HA-EDS1 in Sf9 cells using baculovirus system | This study |

Kindly donated by <sup>1</sup>Sophien Kamoun, <sup>2</sup>Suayib Üstün, <sup>3</sup>Jonathan Jones and <sup>4</sup>Tolga Bozkurt.

**Table S7.**
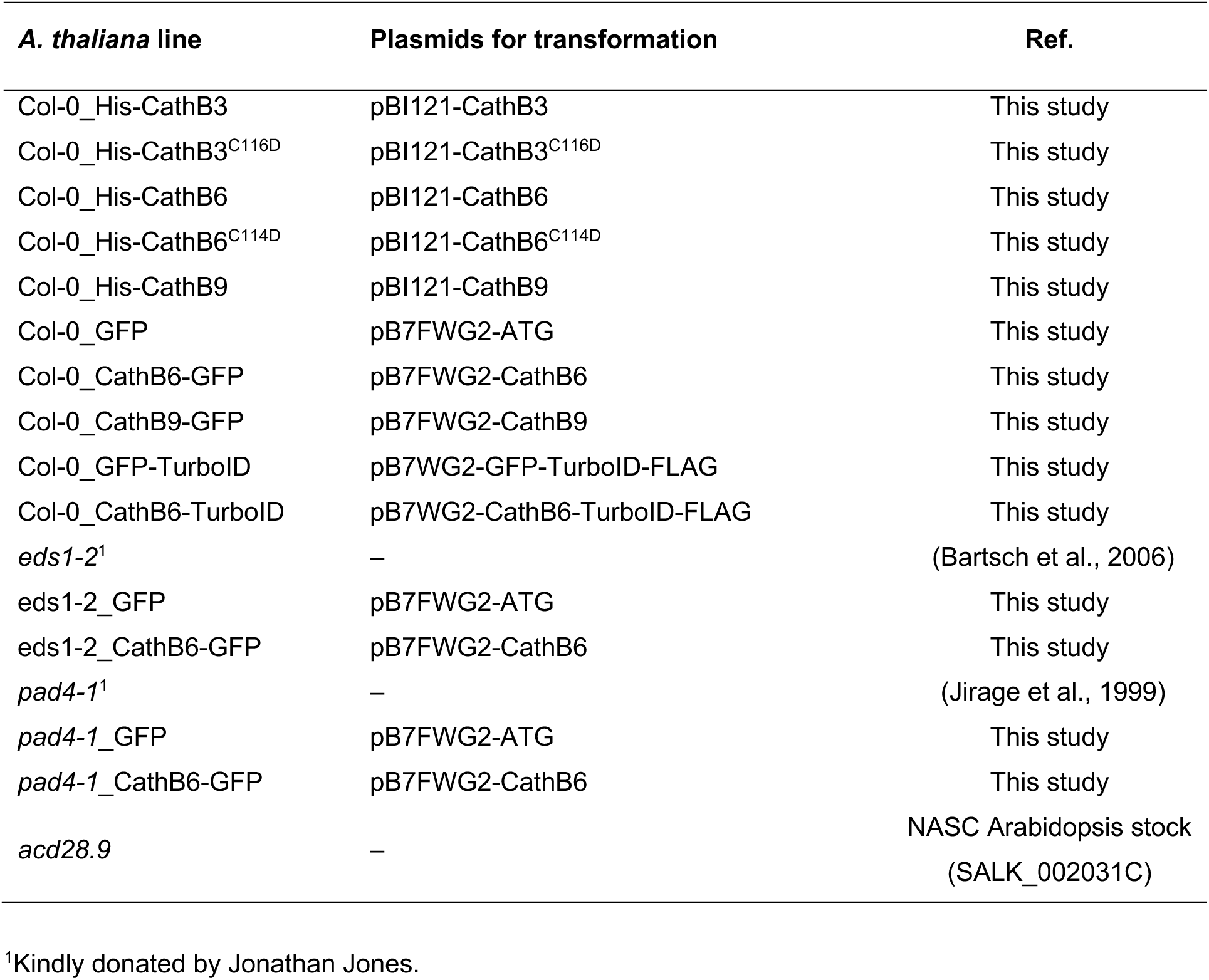
*A. thaliana* lines used in the study.

| ***A. thaliana* line** | **Plasmids for transformation** | **Ref.** |
| --- | --- | --- |
| Col-0_His-CathB3 | pBI121-CathB3 | This study |
| Col-0_His-CathB3<sup>C116D</sup> | pBI121-CathB3<sup>C116D</sup> | This study |
| Col-0_His-CathB6 | pBI121-CathB6 | This study |
| Col-0_His-CathB6<sup>C114D</sup> | pBI121-CathB6<sup>C114D</sup> | This study |
| Col-0_His-CathB9 | pBI121-CathB9 | This study |
| Col-0_GFP | pB7FWG2-ATG | This study |
| Col-0_CathB6-GFP | pB7FWG2-CathB6 | This study |
| Col-0_CathB9-GFP | pB7FWG2-CathB9 | This study |
| Col-0_GFP-TurboID | pB7WG2-GFP-TurboID-FLAG | This study |
| Col-0_CathB6-TurboID | pB7WG2-CathB6-TurboID-FLAG | This study |
| *eds1-2*<sup>1</sup> | – | (Bartsch et al., 2006) |
| *eds1-2_GFP* | pB7FWG2-ATG | This study |
| *eds1-2_CathB6-GFP* | pB7FWG2-CathB6 | This study |
| *pad4-1*<sup>1</sup> | – | (Jirage et al., 1999) |
| *pad4-1_GFP* | pB7FWG2-ATG | This study |
| *pad4-1_CathB6-GFP* | pB7FWG2-CathB6 | This study |
| *acd28.9* | – | NASC Arabidopsis stock<br>(SALK_002031C) |

<sup>1</sup>Kindly donated by Jonathan Jones.

**Movie S1.** *M. persicae* CathB6 locates to mobile puncta within cells of *N. benthamiana* leaves. The white arrow shows an example of a puncta fusion. Scale bar, 30 μm.

